# Antitoxin-induced auto-phosphorylation neutralizes the nucleotidyltransferase toxin AbiEii from *Streptococcus agalactiae* to safeguard global translation

**DOI:** 10.64898/2026.01.19.700257

**Authors:** Tom J. Arrowsmith, Xibing Xu, Sam C. Went, Xue Han, Abbie Kelly, Paul Chansigaud, Zara J. Emsley, Jeremy Dubrulle, Peter C. Fineran, Pierre Genevaux, Tim R. Blower

## Abstract

Nucleotidyltransferases (NTases) control diverse physiological processes, including DNA replication and repair, antibiotic resistance, and RNA modification. The *Streptococcus agalactiae* abortive infection family toxin, AbiEii, belongs to the DUF1814 superfamily of NTases and blocks bacterial growth through an as yet unknown mechanism. Antitoxin AbiEi has been reported to antagonize AbiEii as part of a Type IV toxin-antitoxin system. Here, we demonstrate through structural, biochemical, and biophysical studies that AbiEi binds to AbiEii to form a stable toxin-antitoxin complex in solution. Antitoxin binding ultimately leads to AbiEii auto-phosphorylation and an abolition of NTase activity. These data reclassify AbiE systems as Type VII toxin-antitoxin systems. Toxin phosphorylation appears to be a dynamic and reversible process, and we show that the sole serine/threonine phosphatase *Sa*STP of *S. agalactiae* can efficiently dephosphorylate AbiEii *in vitro*. Mutagenesis studies and functional assays indicate that a widespread mechanism of toxin auto-phosphorylation accounts for AbiEii neutralization, further supporting recent hypotheses outlining control of homologous MenAT toxin-antitoxin systems from *Mycobacterium tuberculosis*.

## Introduction

Since their discovery as plasmid addiction modules, the number and range of cellular functions associated with bacterial toxin-antitoxin (TA) systems has grown to include stress responses, pathogenicity, and bacteriophage (phage) defence^1–4^. One mechanism employed by bacteria as a means of protection against phages is abortive infection (Abi), wherein infected cells suffer irreversible damage as a direct consequence of the initial infection, ultimately leading to cell death that restricts the release of viral progeny and halts the spread of infection^2^. Examples of TA systems shown to function through an Abi mechanism include ToxIN^5^, DarTG^6^, and AbiE^7^, which is the fourth most abundant phage defence system in nature^8^. TA systems such as AbiE are encoded chromosomally or on mobile genetic elements^9,10^, and are conventionally small bicistronic loci, though recent studies have shown not all TA systems fall within this paradigm^11,12^. The toxic component of the TA module typically targets an essential pathway involved in growth or metabolism, such as DNA replication or protein synthesis^13,14^. Toxin activation can be achieved through proteolytic degradation of free labile antitoxins, dysregulation of host transcriptional machinery, or conditional remodeling of stable TA complexes to liberate free active toxin^15–18^, though specific activating triggers remain elusive for most systems. The deleterious activity of the toxin is counteracted by the action of a cognate antitoxin, which can block toxicity using an assortment of neutralization mechanisms, including toxin sequestration, target antagonism, or covalent modification of the toxin’s catalytic site^19^.

*Lactococcus lactis* plasmid pNP40 was shown to encode a functional AbiEi-AbiEii pair that conferred resistance against the 936 family of phages through an Abi mechanism^7^. *L. lactis* AbiE interfered with phage Φ712 DNA packaging late in the phage lytic cycle, thereby preventing the production of viable virions even as the infected cells progressed towards death due to phage infection^7,20^. The human pathogen *Streptococcus agalactiae* also encodes a functional AbiE TA system on a highly conserved integrative conjugative element (ICE) found in β-hemolytic streptococci clinical isolates^21^. This system is comprised of an AbiEi antitoxin (*SAG1284*; *abiEi*) and an AbiEii toxin (*SAG1285*; *abiEii*)^20^. Previous functional characterization demonstrated that *S. agalactiae* AbiEi and AbiEii do not interact when co-expressed in *Escherichia coli*, leading to the conclusion that AbiE belongs to the Type IV classification of TA systems, wherein the antitoxin indirectly negates the activity of the toxin through antagonism of the toxin’s cellular target(s)^19,20^. *S. agalactiae* AbiEii was also previously shown to preferentially bind GTP *in vitro* and demonstrated potent growth inhibition when overexpressed in *E. coli*^20^. Cellular toxicity was efficiently blocked by expression of the AbiEi antitoxin^20^, which has also been shown to transcriptionally autoregulate expression of the *abiE* operon through binding of inverted repeats upstream of its transcriptional start site^22^.

*Mycobacterium tuberculosis*, the aetiological agent of human tuberculosis, encodes the MenAT (*<u>M</u>ycobacterial* Abi<u>E</u>-like <u>N</u>ucleotidyltransferase <u>A</u>ntitoxin <u>T</u>oxin) family of TA systems comprising nucleotidyltransferase (NTase) toxins related to AbiEii, and their antagonistic antitoxin partners^14,23,24^. The four MenT toxins of *M. tuberculosis* and AbiEii of *S. agalactiae* all belong to the DUF1814 superfamily of NTases, which also includes *Paracoccidioides brasiliensis* Pb27, *Klebsiella pneumoniae* Aminoglycoside-2”-O-Nucleotidyltransferase (ANT(2”)-Ia), and *Helicobacter pylori* JHP933, each of which have been proposed to contribute to the virulence and pathogenicity of their hosts^25–27^. NTases have been linked to an array of physiological roles in both prokaryotes and eukaryotes, including RNA processing, genomic replication and repair, and antibiotic resistance^26,28,29^. MenT toxicity occurs by transfer of specific nucleotides to the 3′-CCA motif of tRNA target molecules, modifying the acceptor stem and preventing aminoacylation of these tRNAs, the result of which is a depletion of charged tRNAs required for translation. MenT_1_ and MenT_3_ exhibit pyrimidine specificity for NTase activity with preference for CTP over UTP, with MenT_3_ also displaying target specificity for tRNA^Ser^ isoacceptors^30^, whilst MenT_4_ is GTP specific and displays broader target specificity *in vitro*^23^. Modification of target tRNAs inhibits cellular translation, leading to growth arrest in the absence of antitoxins^14,23^. The essentiality of the processes in which NTases play a role has ensured host organisms have developed efficient modes of regulation for these enzymes^31,32^. Accordingly, the MenT toxins of *M. tuberculosis* have been shown to require varied modes of regulation. MenA_1_-MenT_1_ and MenA_3_-MenT_3_ belong to the Type VII classification of TA systems, wherein antitoxins post-translationally modify their cognate toxins^33^. MenA_1_ and MenA_3_ antitoxins bind to MenT_1_ and MenT_3_ toxins respectively to form heteromeric toxin-antitoxin complexes^14,34^, both of which differ in their relative stabilities^34^. Using a combination of structural and computational approaches, we had previously predicted that MenA antitoxin-induced conformational changes within the MenT active site repositions catalytic Thr/Ser residues (for MenT_1_/MenT_3_ respectively) within proximity of (i) the γ-phosphate of bound nucleotide substrates, and (ii) a strictly conserved catalytic aspartate residue responsible for abstraction of the Thr/Ser hydroxyl proton^34^. In turn, proton abstraction is predicted to prime the phosphoacceptor for nucleophilic attack of the highly co-ordinated NTP γ-phosphate, triggering toxin auto-phosphorylation^34^. Phosphorylated toxins are devoid of NTase activity (**Fig. 1A**), owing to steric occlusion and reduced electropositive charge within the recessed NTase catalytic pocket. In contrast, MenT_4_ does not interact with the MenA_4_ antitoxin and is not subject to phosphorylation, indicating that there is a currently unknown alternative mode of antitoxicity utilized by MenA_4_.

**Figure 1.**
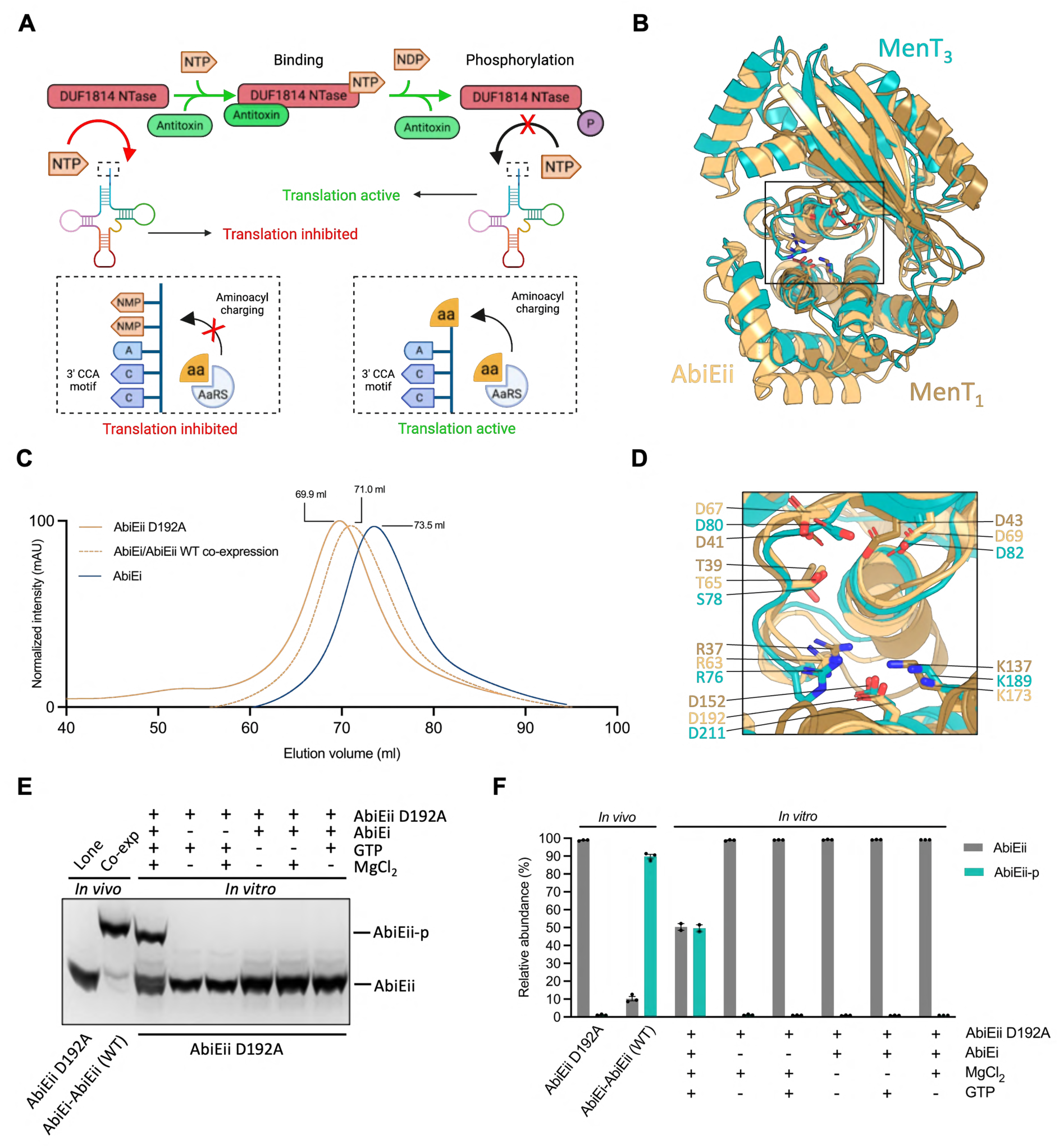
AbiEi-AbiEii constitutes a type VII TA system. (**A**) Schematic diagram of antitoxin-induced auto-phosphorylation for DUF1814 superfamily NTase toxins. In the absence of antitoxin, the NTase toxin transfers nucleotides to the 3′-CCA motif of target tRNAs, preventing aminoacylation (translation inhibited). Antitoxin binding induces a conformational change to the toxin’s active site that repositions the catalytic phosphoacceptor within proximity of (i) a strictly conserved catalytic aspartate responsible for proton abstraction, and (ii) the γ-phosphate of bound nucleotide. Proton abstraction triggers nucleophilic attack of the nucleotide γ-phosphate and results in toxin phosphorylation, blocking NTase activity (translation active). (**B**) Superposition of AbiEii (AlphaFold model; pTM score = 0.92) onto MenT_1_ (PDB 8AN4) and MenT_3_ (PDB 8RR6). Structures are shown as cartoon representations colored wheat (AbiEii), sand (MenT_1_), and teal (MenT_3_). (**C**) Overlay of Size Exclusion Chromatography (SEC) traces corresponding to AbiEii D192A and AbiEi expressed alone, and AbiEii WT expressed in the presence of AbiEi. Samples were analyzed using a HiPrep^™^ 16/60 Sephacryl^®^ S-200 HR column. The AbiEi-AbiEii WT co-expression sample eluted at 71.0 ml, corresponding to a mass matching the calculated molecular weight (M_w_) of AbiEii. Chromatograms are normalized between 0-100 for presentation and comparison, cropped to the appropriate scale. (**D**) Close-up view of the boxed region in (**B**) highlighting strictly conserved toxin active site residues, shown as sticks with atoms colored red for oxygen and blue for nitrogen. (**E**) Phos-Tag SDS-PAGE analysis of either AbiEii D192A expressed alone, AbiEii WT expressed in the presence of AbiEi, or AbiEii D129A incubated with combinations of AbiEi, MgCl_2_, and GTP. (**F**) Densitometric analysis of Phos-Tag SDS-PAGE bands corresponding to AbiEii (WT or D192A mutant) and AbiEii-p from (**E**). Assays shown are representative of three independent biological replicates and plotted data represent the mean ± SEM.

In this study we leveraged knowledge gained from characterization of the MenAT systems to perform biochemical, structural, and functional characterization of *S. agalactiae* AbiE. We have demonstrated, against previous literature, that neutralization of the AbiEii toxin is attributable to interaction with the antitoxin AbiEi, leading to phosphorylation of the toxin. This indicates that AbiE is not a Type IV system, but rather another member of the recently discovered Type VII systems, unveiling a widespread mechanism of inducible auto-phosphorylation amongst species of unrelated pathogenic bacteria. Biochemical analyses show that, like the recently characterized MenT_1_ and MenT_3_ of toxins of *M. tuberculosis,* AbiEii is phosphorylated at the structurally homologous phosphoacceptor residue T65. Binding assays and crystallographic studies reveal that AbiEi and AbiEii form a heterodimeric complex in solution, whilst functional characterization highlights key AbiEi residues directly involved in neutralization of AbiEii toxicity. Autoradiography experiments demonstrate that the toxin exhibits nucleotide specificity for UTP, highlighting additional variability in the substrate specificities of related DUF1814 NTases.

## Results

### AbiEi-AbiEii constitutes a Type VII TA system

It was previously reported that *S. agalactiae* AbiEi-AbiEii belongs to the Type IV classification of TA systems, owing to the inability of the AbiEii toxin to pull-down its cognate AbiEi antitoxin in bait-prey co-precipitation experiments^20^. *S. agalactiae* AbiEii shares structural homology with the related DUF1814 MenT toxins from *M. tuberculosis*. Despite only sharing 16.3% and 16.9% sequence identity with MenT_1_ and MenT_3_, respectively, sequence-independent structural alignment (superposition) of AbiEii (AlphaFold model; pTM score = 0.92) onto the solved crystal structures of MenT_1_ (PDB 8AN4) and MenT_3_ (PDB 8RR6) returned RMSD values of 4.839 Å (across 964 atoms) and 2.227 Å (across 1166 atoms), again respectively, reflecting shared structural homology between these NTases (**Fig. 1B**). Furthermore, superposition of related COG5340 antitoxins AbiEi (PDB 6Y8Q^35^) and MenA_3_ (AlphaFold model; pTM score = 0.87) returned an RMSD of 3.787 Å (across 1517 atoms) (**Supplementary Fig. S1A**). We therefore hypothesized that AbiE is not a Type IV TA system and might use a similar mode of phosphorylation-dependent antitoxicity against the cognate AbiEii toxin as that used by MenA_3_ against MenT_3_^14^. We first attempted to express AbiEii in the absence or presence of AbiEi, though lone expression of the wild-type (WT) toxin was highly refractory to *E. coli* growth, precluding purification and downstream analysis. To bypass the intoxicating effects of AbiEii WT when expressed in the absence of antitoxin, we utilized a non-toxic alanine substitution mutant, D192A^20^, as a surrogate for lone toxin expression. Using this inactive catalytic mutant, we were able to purify sufficiently high yields of the toxin to allow for comparison of lone toxin, lone antitoxin, and toxin-antitoxin co-expression samples using Size Exclusion Chromatography (SEC). In agreement with existing literature^20^, co-expression of AbiEii WT and AbiEi failed to result in a substantial reduction in elution volume during SEC relative to when AbiEii D192A was expressed alone (**Fig. 1C**), whilst SDS-PAGE analysis of peak chromatographic elution fractions also confirmed an absence of AbiEi (**Supplementary Fig. S1B**). These data show that AbiEii and AbiEi do not form a stable complex when co-overexpressed in *E. coli*.

To establish whether AbiEii was phosphorylated in the presence of AbiEi, we analyzed the purified co-expression sample using positive electrospray ionization time-of-flight mass spectrometry (Es^+^-ToF MS) and liquid chromatography tandem mass spectrometry (LC-MS/MS). Es^+^-ToF MS revealed the mass of AbiEii co-expressed in the presence of AbiEi to be 33318 Da (**Supplementary Fig. S1C**), 80 Da higher than the expected mass of the native hexahistidine-tagged protein and matching the expected mass of a single phosphate group. Furthermore, LC-MS/MS analysis identified the presence of a phosphothreonine at position T65 (**Supplementary Fig. S1D**), confirming the presence of phosphorylated AbiEii (herein referred to as AbiEii-p) following co-expression with AbiEi. Close-up views of the AbiEii AlphaFold model superposed onto MenT_1_ and MenT_3_ reveals that T65 is likely the structurally equivalent residue to the T39 and S78 phosphoacceptors of MenT_1_ and MenT_3_, respectively (**Fig. 1D**)^14,34^. Therefore, *S. agalactiae* AbiEi-AbiEii likely constitutes a Type VII TA system, and not a Type IV system as previously reported^20^.

Previous work identified Mg^2+^ and nucleotide substrates as essential components for MenT_1_ phosphorylation, whilst also demonstrating that the non-toxic MenT_1_ D152A mutant remains competent for phosphorylation, albeit to a lesser degree than its WT counterpart^34^. AbiEii D192 is structurally equivalent to MenT_1_ D152, and so we reasoned that the AbiEii D192A mutant may also remain competent for phosphorylation. We applied the same methodology to the AbiEi-AbiEii TA system to characterize the essential components for phosphorylation, by incubating AbiEii D192A with combinations of MgCl_2_, AbiEi, and GTP, reportedly the preferred NTase substrate of AbiEii^20^. Samples were then analyzed by Phos-Tag SDS-PAGE and Es^+^-ToF MS. We observed that *in vivo* co-expression of WT AbiE proteins yielded a majority of AbiEii-p (**Figs. 1E and F**). In contrast, *in vitro* co-incubation of AbiEii D192A with AbiEi, MgCl_2_, and GTP generated a heterogeneous population of AbiEii D192A and AbiEii-p D192A of roughly equal abundancies (**Figs. 1E and F**), though the comparably lower levels of AbiEii-p could be attributable to weaker phosphorylation activity of the D192A mutant compared to AbiEii WT when co-expressed. Akin to MenT_1_, no phosphorylation was detected when AbiEii D192A was incubated in the absence of either AbiEi, MgCl_2_, or NTP (**Figs. 1E and F**; **Supplementary Fig. S1E**). Es^+^-ToF MS also revealed that AbiEii D192A was phosphorylated when incubated with MgCl_2_, AbiEi, and either ATP, GTP, CTP, or UTP (**Supplementary Fig. S1F**), with an absence of phosphorylation when MgCl_2_ was omitted from each reaction (**Supplementary Fig. S1G**). These data indicate that AbiEii likely exhibits different nucleotide specificities for NTase and phosphorylation activities and, like MenT_1_, can utilize all nucleotides as substrates for phosphorylation.

### Phosphorylation of AbiEii is reversible

Under favorable physiological growth conditions, AbiEii likely exists in its phosphorylated form, as evidenced by the production of a homogenous population of AbiEii-p when co-overexpressed with AbiEi in *E. coli* (**Figs. 1E and F**). Whilst a homogeneous population of AbiEii-p might confer a survival advantage to *S. agalactiae*, ensuring the toxin remains inert in the absence of stress, existing stockpiles of phosphorylated toxin would be functionally redundant if the cell were to encounter stressful stimuli or specific activating triggers. Having to first transcribe and translate new unphosphorylated AbiEii would cause the pathogen to be slower in deploying evasive countermeasures in a bid to arrest growth. We therefore reasoned that cellular phosphatases might play a role in toxin reactivation, providing the host with a more rapid means of regenerating active toxin than *de novo* synthesis. It was previously shown that the *M. tuberculosis* serine/threonine phosphatase PstP (*rv0018c*) could non-specifically dephosphorylate MenT_3_-p *in vitro*^36^, though restoration of NTase activity was never explored. We identified a homologue of PstP from *S. agalactiae* sharing 33% sequence identity with PstP (*stp1*; herein referred to as *Sa*STP), with high conservation of putative active site residues proposed to contribute to phosphatase activity^37^ (**Fig. 2A**). Superposition of respective homologs returned an RMSD of 1.257 Å (across 957 atoms), reflective of high structural homology, whilst close-up views highlighted remarkable co-localization of active site residues (**Figs. 2B and C**).

**Figure 2.**
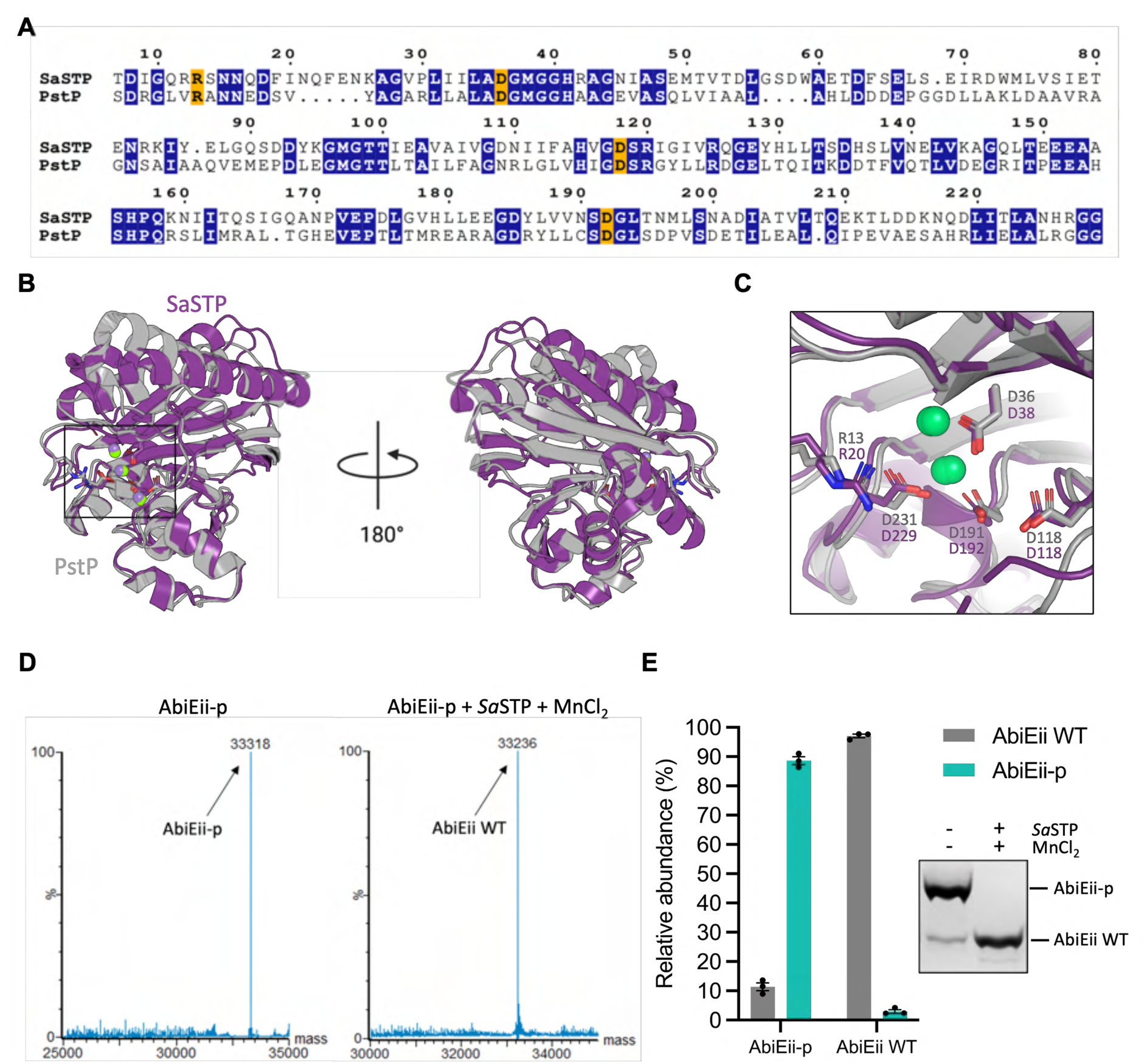
*Sa*STP fully dephosphorylates AbiEii-p *in vitro*. (**A**) Pairwise sequence alignment of *S. agalactiae Sa*STP and *M. tuberculosis* PstP. Strictly conserved residues are depicted by bold white text either shaded blue (non-catalytic) or orange (catalytic). (**B**) Superposition of *S. agalactiae Sa*STP (PDB 2PK0) and *M. tuberculosis* PstP (PDB 1TXO), with front and rear views shown. Structures are shown as cartoon representations colored purple (*Sa*STP) and grey (PstP). Mn^2+^ ions are shown as spheres and are colored lime green. (**C**) Close-up view of the boxed region in (**B**). Conserved putative active site residues are shown as sticks with atoms colored red for oxygen and blue for nitrogen. (**D**) Es^+^-ToF MS analysis of AbiEii-p incubated in the absence and presence of *Sa*STP and MnCl_2_. (**E**) Phos-Tag SDS-PAGE and densitometric analysis following incubation of AbiEii-p in the absence and presence of *Sa*STP and MnCl_2_. Assays shown are representative of three independent biological replicates and plotted data represent the mean ± SEM.

To test whether AbiEii-p could be dephosphorylated *in vitro*, we expressed and purified *Sa*STP (**Supplementary Figs. S2A and B**), then subsequently assessed its ability to dephosphorylate AbiEii-p in phosphatase assays. Co-incubation of AbiEii-p and *Sa*STP in the presence of MnCl_2_ resulted in an 80 Da reduction in mass relative to toxin incubated alone (**Fig. 2D**). Phos-Tag SDS-PAGE confirmed this reduction in mass to be the result of dephosphorylation (**Fig. 2E**). We subsequently repeated *in vitro* dephosphorylation reactions on a larger scale and isolated pure, non-phosphorylated AbiEii (AbiEii WT) by using nickel resin to sequester the hexahistidine-tagged toxin (**Supplementary Fig. S2C**). This allowed us to circumvent the extreme toxicity of AbiEii in *E. coli* and generate non-phosphorylated toxin for biochemical and crystallographic studies.

Having now successfully isolated AbiEii WT, we assessed the ability of the AbiEi antitoxin to induce a second round of phosphorylation by co-incubating the respective proteins in the presence of GTP and MgCl_2_. Co-incubation resulted in an increase in the mass of AbiEii WT by 82 Da relative to when incubated in the absence of antitoxin (**Supplementary Fig. S2D**), confirming that dephosphorylation restores competency for subsequent phosphorylation events. Collectively, these data show that the phosphorylation status of AbiEii can be reversibly and antagonistically manipulated *in vitro* by *Sa*STP and AbiEi. The exact role of cellular phosphatases in toxin re-activation *in vivo* remains to be explored.

### AbiEi-AbiEii form a stable heterodimer in solution

Our data indicated that AbiEii is phosphorylated when co-expressed with AbiEi (**Supplementary Fig. S1C**), and that phosphorylation is dependent on AbiEi (**Figs. 1E and F**). These findings led us to question the previous conclusion that AbiEi and AbiEii do not interact, as (i) both components likely interact to allow phosphorylation, and (ii) previous investigations used co-expressed and thereby phosphorylated AbiEii, which appears reduced in ability to bind AbiEi^20^. Accordingly, we co-incubated AbiEii-p or AbiEii WT with AbiEi, then analyzed the resulting mixtures by SEC. Co-incubation of AbiEii-p and AbiEi failed to reduce column elution volume (V_e_) relative to when the toxin was incubated alone (**Fig. 3A**), corroborating the lack of an observable complex co-precipitated from *in vivo* samples when the toxin is phosphorylated^20^. In contrast, co-incubation of AbiEii WT and AbiEi resulted in an ∼15.3 ml reduction in column V_e_ compared to when the toxin was incubated alone (**Fig. 3A**), reflecting the formation of a larger complex in solution. SDS-PAGE analysis of peak chromatographic fractions corresponding to each major peak observed during SEC confirmed the presence of AbiEi in the AbiEii WT + AbiEi co-incubation peak, but not that of the AbiEii-p+ AbiEi mixture (**Fig. 3B**), confirming the observed reduction in V_e_ to be the result of stable TA complex formation.

**Figure 3.**
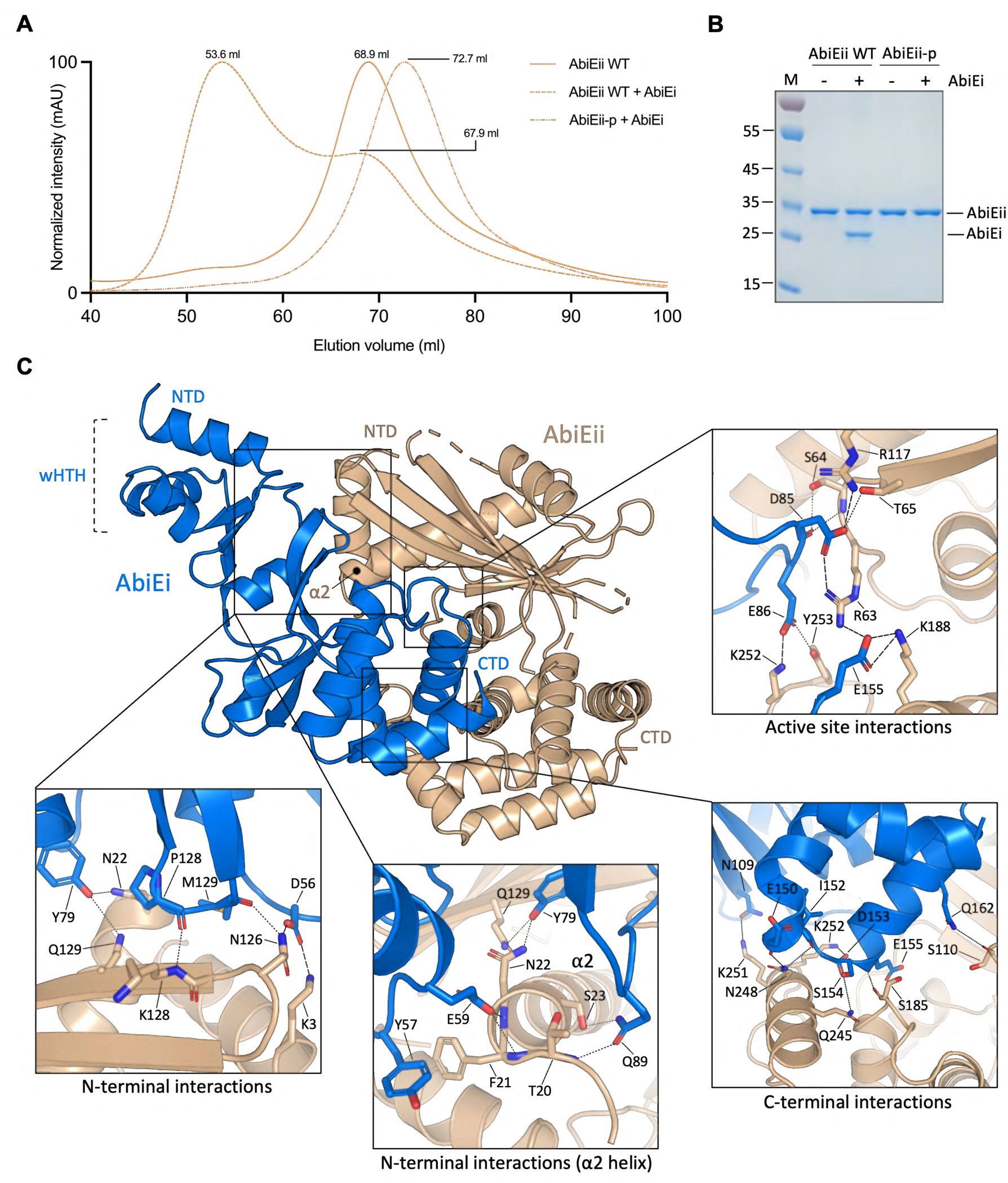
The AbiEi C-terminal antitoxicity domain forms a network of interactions with AbiEii. (**A**) Overlaid SEC traces corresponding to AbiEii WT incubated alone, and either AbiEii WT or AbiEii-p incubated in the presence of AbiEi. Samples were analyzed using a HiPrep^™^ 16/60 Sephacryl^®^ S-200 HR column. Chromatograms are representative of three independent biological replicates and are normalized between 0-100 for presentation and comparison, cropped to the appropriate scale. (**B**) SDS-PAGE analysis of peak chromatographic fractions from (**A**) corresponding to lone toxin or toxin-antitoxin incubations. (**C**) Crystal structure of the AbiEi:AbiEii heterodimer (PDB 9HLO) shown as a cartoon representation colored marine (AbiEi) and wheat (AbiEii), with close-up views highlighting interaction networks located at the AbiEii N-terminus, C-terminus, and active site. Residues are shown as sticks with atoms colored red for oxygen and blue for nitrogen. Interaction types are denoted by lines; dotted line = hydrogen bond, dashed line = salt bridge.

Having demonstrated that AbiEi can bind to AbiEii WT, we repeated binding assays using an analytical Superdex™ 75 increase 10/300 GL SEC column to help in determining the stoichiometry of the TA complex observed during SEC (**Fig. 3A; Supplementary Fig. S3A**). In the absence of AbiEi, AbiEii WT eluted from the SEC column at 12.90 ml (**Supplementary Fig. S3A**). However, in the presence of antitoxin, the major species eluted ∼1.9 ml earlier, with SDS-PAGE analysis of the co-incubation mixture once again confirming pull-down of AbiEi (**Supplementary Fig. S3A**). Correlation of the observed molecular weight (M_w_) for this species against the calculated M_w_ of AbiEii returned a ratio of 1.93, suggesting the complex to be approximately twice the expected size of the AbiEii toxin and alluding to the existence of an AbiEi:AbiEii heterodimer in solution. Based on these observations, we generated a predictive model of the theoretical AbiEi:AbiEii heterodimer using AlphaFold (pTM score = 0.79) to allow for comparison of observed M_w_ and hydrodynamic radii (R_st_) against that of the predictive model. The observed V_e_ of the unknown species captured during SEC (10.99 ml) closely matches the calculated V_e_ of the theoretical AbiEi:AbiEii heterodimer at ∼11.7 ml (**Supplementary Fig. S3A**), whilst correlation of observed and calculated M_w_ and R_st_ values returned ratios of 1.14 and 1.10 respectively (**Supplementary Fig. S3B**), indicating that the species captured by SEC is likely that of an AbiEi:AbiEii heterodimer. To confirm that AbiEii D192A can also form a stable TA complex and ensure that previous assays utilizing this non-toxic mutant were representative, we repeated incubations *in vitro* and analyzed the resulting mixtures by mass photometry. In agreement with analytical SEC, mass photometry analysis of AbiEii D192A incubated with or without AbiEi supported the existence of a heterodimeric complex in solution following co-incubation of AbiEii D192A and AbiEi (**Supplementary Fig. S3C**), with the observed mass of the species closely matching the calculated mass of the AbiEi:AbiEii heterodimer at ∼56 kDa.

The isolated AbiE WT TA complex captured during SEC was concentrated and put into crystallographic trials. This produced thick cuboidal crystals in condition screens containing 0.2 M potassium acetate, 0.1 M MES pH 6.0, and 15% v/v pentaerythritol ethoxylate. Crystals were subsequently harvested and analyzed using X-ray diffraction. Using these data, the structure of the AbiEi:AbiEii heterodimer was determined to a resolution of 2.2 Å (**Fig. 3C and Table 1**). Within the resolved structure, the AbiEii toxin adopts the expected NTase-like fold comprising central β-sheets flanked by alternating α-helices. Four unresolved gaps are also present in external flexible loops within the N-terminal domain of AbiEii between positions 16-17, 91-95, 111-113, and 138-143 inclusively. PISA analysis^38^ identified three primary interfaces between respective proteins, localized to the toxin’s N-terminal domain (NTD), C-terminal domain (CTD), and the conserved active site.

**Table 1.** Data collection and refinement statistics.

|  | AbiEi:AbiEii |
| --- | --- |
| PDB ID Code | 9HLO |
| **Data collection** |  |
| Space group | I 1 2 1 |
| Cell dimensions |  |
| *a*, *b*, *c* (Å) | 84.355 86.796 82.245 |
| $\alpha$, $\beta$, $\gamma$ (°) | 90 118.266 90 |
| Resolution (Å) | 72.78 - 2.20 (2.28 - 2.20) |
| *R*<sub>merge</sub> | 0.014 (0.074) |
| *R*<sub>meas</sub> | 0.020 (0.105) |
| *I* / *σI* | 26.2 (3.9) |
| Completeness (%) | 99.99 (100.00) |
| Redundancy | 2.0 (2.0) |
| **Refinement** |  |
| Resolution (Å) | 2.20 |
| No. reflections | 54092 (4689) |
| Unique reflections | 27558 (2757) |
| *R*<sub>work</sub> | 0.2278 (0.2629) |
| *R*<sub>free</sub> | 0.2727 (0.3395) |
| No. atoms | 3937 |
| Protein | 3805 |
| Ligand/ion | 21 |
| Water | 111 |
| *B*-factors | 46.31 |
| Protein | 46.38 |
| Ligand/ion | 54.70 |
| Water | 42.37 |
| R.m.s. deviations |  |
| Bond lengths (Å) | 0.008 |
| Bond angles (°) | 0.94 |

Values in parentheses are for highest-resolution shell.

AbiEi forms an extensive network of hydrogen bonds with a handful of AbiEii NTD residues. AbiEi D56 forms a salt bridge and hydrogen bond with AbiEii K3 and N126, respectively, the latter of which also hydrogen bonds with the main-chain carbonyl of AbiEi M129 (**Fig. 3C**). These interactions are supported by hydrogen bonds between the backbone carbonyl of AbiEi P128 and the main-chain amide of AbiEii K128. Several other hydrogen bonds are formed between AbiEi and the AbiEii α2 helix. The side-chain of AbiEi Q89 hydrogen bonds with the main-chain amide nitrogen of AbiEii T20 and the side-chain hydroxyl of AbiEii S23, whilst AbiEi E59 interacts with the backbone amide nitrogen atoms of AbiEii residues F21 and N22 (**Fig. 3C**). These interactions are further stabilized by base-stacking between AbiEii F21 and AbiEi Y57, and hydrogen bonding between AbiEii N22 and the tyrosine hydroxyl of AbiEi Y79, which also interacts with AbiEii Q129. At the toxin’s C-terminus, K251 and Q245 hydrogen bond with AbiEi N109 and S154 respectively (**Fig. 3C**), whilst K252 forms a salt bridge with the carboxylate side-chain of AbiEi D153. Toxin residues S185 and N248 form hydrogen bonds with the main-chain oxygen and nitrogen atoms of AbiEi E155 and E150/I152, respectively. These interactions and further supported by a final hydrogen bond between AbiEii S110 and AbiEi Q162.

Within the AbiEii active site, the flexible antitoxin loop connecting α6-β4 protrudes into the nucleotide-binding pocket and remains anchored by a network of salt bridges and hydrogen bonds with catalytically important toxin residues. The carboxylate side-chain of AbiEi D85 forms salt bridges with AbiEii R117 and R63, whilst AbiEi E86 forms a salt bridge with AbiEii K252 (**Fig. 3C**). AbiEi D85 and E86 also form hydrogen bonds with AbiEii; D85 directly interacts with the T65 phosphoacceptor and adjacent S64 residue via side-chain and main-chain contacts, respectively, whilst E86 contacts the C-4 hydroxyl of Y253. In addition to the interaction with AbiEi D85, AbiEii R63 forms a second salt bridge with AbiEi E155, which itself forms two salt bridges with AbiEii K188. This extensive matrix of ionic interactions ensures that R63 is pulled away from the nucleotide-binding pocket of the toxin. Crucially, R63 was previously shown to be essential to AbiEii toxicity^20^, and structurally equivalent residues in other DUF1814 homologs co-ordinate the γ-phosphate of incoming nucleotide substrates^26^.

Despite the extensive network of hydrogen bonds and salt bridges formed between respective proteins, very few contacts are made between AbiEii and the winged helix-turn-helix DNA-binding domain at the N-terminus of AbiEi (**Fig. 3C**), supporting previous work showing antitoxic activity is confined to the antitoxin’s CTD^20^. Meanwhile, DNA-binding residues in both the AbiEi NTD and CTD remain exposed in the resolved heterodimer (**Supplementary Fig. S3D**), corroborating previous data showing that co-induction of AbiEii has no effect on *abiEi-abiEii* autoregulation by AbiEi^20^. Binding of the AbiEi CTD across the electropositive face of AbiEii also results in steric occlusion of the toxin’s active site that might prevent access to incoming tRNA targets (**Supplementary Fig. S3D**), potentially hinting at binding as a viable mode of antitoxicity in the absence of phosphorylation. These structural data provide the first experimental detail of an AbiE TA complex and reclassify AbiE TA systems from Type IV to Type VII.

### AbiEii phosphorylation inhibits nucleotidyltransferase activity

Next, we examined the functional impact of antitoxin binding and toxin phosphorylation on NTase activity, as structural data had indicated that binding alone may be sufficient for neutralization of AbiEii. First, the activity of AbiEii WT and AbiEii-p were compared to ascertain whether phosphorylation impacts NTase activity, as had been observed for MenT_1_ and MenT_3_^14,34^, and whether dephosphorylation could restore NTase activity. Five μM of purified AbiEii WT or AbiEii-p were added to cell-free transcription/translation protein synthesis reactions (PURExpress^®^; NEB) to quantify changes in DHFR control protein expression. AbiEii WT completely blocked DHFR expression *in vitro*, whilst AbiEii-p failed to inhibit NTase activity (**Fig. 4A**), demonstrating that phosphorylation blocks NTase activity, and confirming dephosphorylation successfully restores NTase activity.

**Figure 4.**
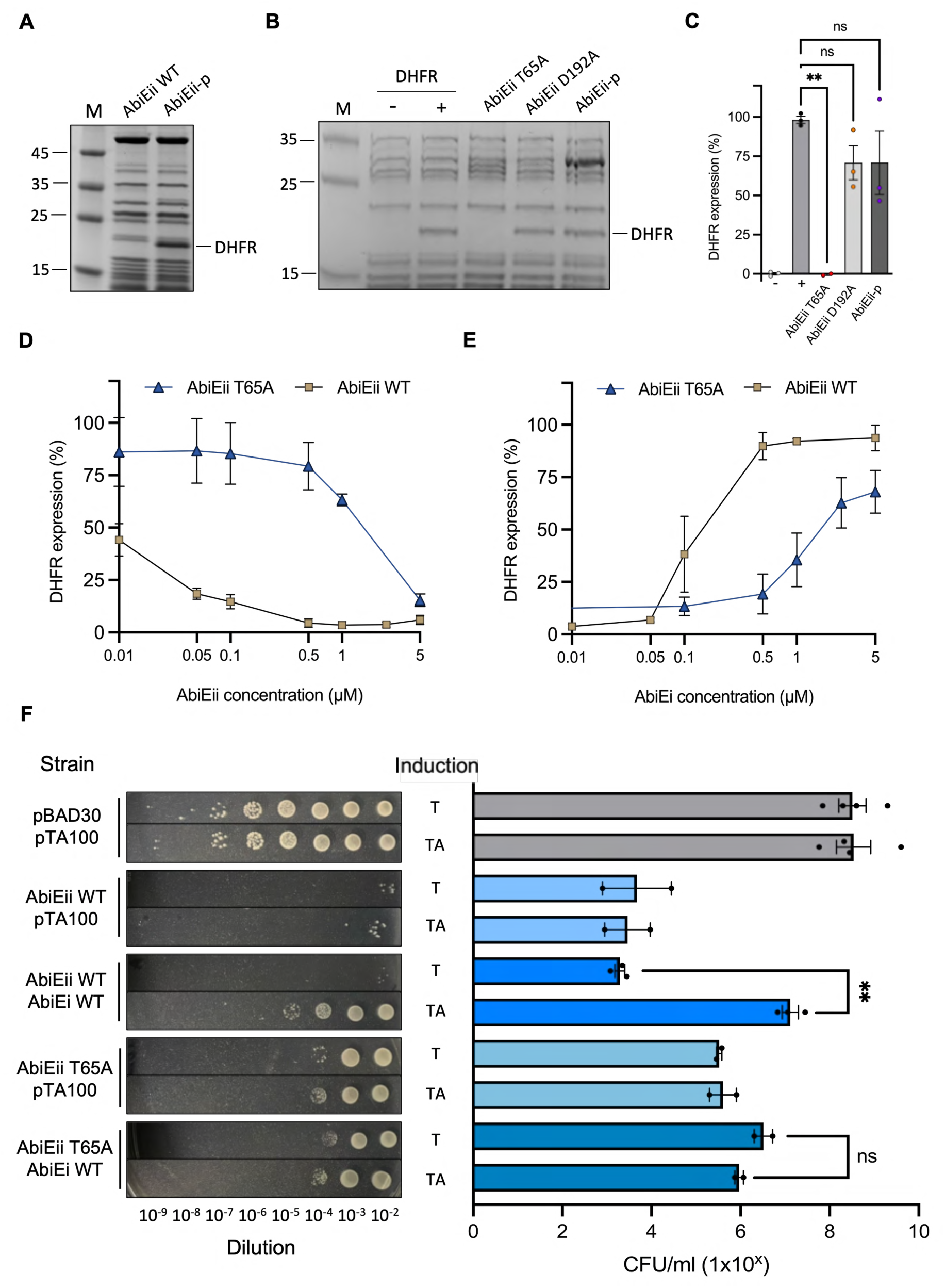
T65 is important, but not essential, for NTase and phosphorylation activity. (**A**) *In vitro* transcription/translation reactions assessing levels of DHFR control protein produced in the presence of either 5 µM AbiEii WT or AbiEii-p. (**B**) *In vitro* transcription/translation reactions assessing levels of DHFR control protein produced in the absence and presence of 5 µM AbiEii T65A, AbiEii D192A, or AbiEii-p. Reactions were loaded onto 4-20% gradient gels ran at 180 V for 1 h. (**C**) Densitometric analysis of SDS-PAGE bands from (**B**) corresponding to DHFR. (**D**) Dose-response curves of increasing concentrations of either AbiEii WT or AbiEii T65A plotted against DHFR expression, normalized against DHFR expressed in the absence of toxin (100% expression). Data points were generated following densitometric analysis of bands corresponding to DHFR from **Supplementary Figs. S4A and B**. (**E**) Logarithmic dose-response curves of either 5 µM AbiEii WT or AbiEii T65A incubated with increasing concentrations of AbiEi plotted against DHFR expression, normalized against DHFR expressed in the absence of toxin (100% expression). Data points were generated following densitometric analysis of bands corresponding to DHFR from **Supplementary Figs. S4C and D**. Logarithmic curves were generated using non-linear regression in Prism (GraphPad). (**F**) Endpoint viable count antitoxicity assays of *E. coli* DH5α transformed with either pBAD30 empty vector, pTRB482 (AbiEii WT), or pTRB736 (AbiEii T65A), and either pTA100 empty vector or pTRB481 (AbiEi). Overnight cultures were re-seeded into fresh LB supplemented with Ap, Sp and D-glu and grown to mid-log phase. Samples were serially diluted 10^−2^-10^−9^ and spotted onto M9A plates containing Ap and Sp, with or without D-glu, L-ara, and IPTG for repression of toxin expression, induction of toxin expression, and induction of antitoxin expression, respectively. Plates were incubated at 37 °C for 48 h, after which they were imaged and colonies counted to determine CFU/ml (one-way ANOVA, P < 0.05). “T” = toxin induced, “TA” = antitoxin + toxin induced. Assays shown are representative of three independent biological replicates and plotted data represent the mean ± SEM.

Previous work showed that the MenT_1_ phosphoacceptor T39 plays an important role in the catalytic cycle of the enzyme, but is not essential for toxicity in *M. smegmatis*^34^. Similarly, the phosphorylation-deficient MenT_1_ T39A mutant could be blocked by MenA_1_ through binding alone *in vitro*, albeit less efficiently than MenT_1_ WT, where MenA_1_ binding would lead to phosphorylation. This indicated that T39 was not essential for antitoxicity, which was subsequently verified by the ability of MenA_1_ to block MenT_1_ T39A toxicity in *M. smegmatis*^34^. To quantify the functional importance of AbiEii T65 in modes of toxicity and antitoxicity, we first expressed and purified AbiEii T65A in the absence of antitoxin, then added the toxin to PURExpress^®^ reactions and measured its ability to block protein synthesis. When added at a final concentration of 5 μM, AbiEii T65A efficiently blocked DHFR expression (one-way ANOVA, P = 0.0020), whilst AbiEii D192A and AbiEii-p had little effect on protein synthesis (one-way ANOVA, P = 0.45; **Figs. 4B and C**). This result indicated that AbiEii T65A was still functional as an NTase and corroborated the reported lack of AbiEii D192A toxicity *in vivo*^20^. Reactions were then repeated by titrating AbiEii WT and T65A mutant to quantify the relative ability of each to inhibit DHFR expression across a range of concentrations. Both AbiEii WT and T65A inhibited DHFR expression in a concentration-dependent manner, with AbiEii T65A capable of blocking protein synthesis by >75% when added to PURExpress^®^ reactions at 5 μM final concentration (**Fig. 4D and Supplementary Fig. S4A**). In contrast, AbiEii WT achieved comparable levels of protein synthesis inhibition when added to reactions at 0.1 μM final concentration (**Fig. 4D and Supplementary Fig. S4B**). This confirmed that T65 plays an important, but not essential, role in NTase activity.

The dependence of AbiEii phosphorylation on an interaction with the cognate AbiEi antitoxin (**Figs. 1E and F**), coupled with structural evidence for steric occlusion of the toxin’s active site (**Fig. 3D**), led us to speculate whether binding alone may partially block AbiEii NTase activity. To characterize the role of T65 in antitoxicity, we first measured inhibition of AbiEii WT and T65A by the AbiEi antitoxin using PURExpress^®^ assays by titrating increasing concentrations of AbiEi against 5 μM of either toxin variant. AbiEi could partially block AbiEii WT and T65A activity when added to protein synthesis reactions at final concentrations ≤ 0.1 μM (**Fig. 4E**; **Supplementary Figs. S4C and D**). However, above 0.1 μM, AbiEi blocked the activity of AbiEii WT far more efficiently than AbiEii T65A (**Fig. 4E**; **Supplementary Figs. S4C and D**), suggesting that phosphorylation is a more effective means of toxin inhibition than binding alone. Together these data demonstrate that, *in vitro*, AbiEii T65A can be blocked by AbiEi binding, but phosphorylation leads to far more potent AbiEii inhibition.

Finally, we tested whether AbiEii T65A could also be blocked by AbiEi *in vivo* by transforming *E. coli* with either pBAD30 empty vector, AbiEii WT, or AbiEii T65A, and either pTA100 empty vector or AbiEi. Induction of AbiEii WT alone resulted in potent growth inhibition relative to the empty vector control strain (**Fig. 4F**), whilst co-induction of AbiEi efficiently blocked AbiEii WT toxicity (one-way ANOVA, P = 0.0023). Induction of AbiEii T65A alone generated an intermediate phenotype whereby toxicity was reduced by two orders of magnitude compared to AbiEii WT (**Fig. 4F**). However, co-induction of AbiEi had no effect on the number of viable colonies formed (one-way ANOVA, P = 0.7815), indicating that phosphorylation is the sole mode of antitoxicity *in vivo*, and that neutralization of AbiEii T65A *in vitro* could be the result of higher protein concentration than present in cells.

### Structural alignments support antitoxin-induced AbiEii auto-phosphorylation

Despite having shown that AbiEii phosphorylation is dependent on AbiEi (**Figs. 1E and F**), it remained unclear whether AbiEi acts as a protein kinase, or whether the antitoxin induces toxin auto-phosphorylation akin to the *M. tuberculosis* MenA_1_-MenT_1_ and MenA_3_-MenT_3_ systems^34^. If AbiEi were to directly act as a kinase, it would be expected to bind nucleotide substrate(s). To investigate nucleotide binding, we incubated AbiEi in the absence and presence of each nucleotide, then measured the thermal stability of the antitoxin by thermal shift assay. No significant differences in thermal stability were detected between any of the four antitoxin-nucleotide mixtures relative to AbiEi incubated alone (one-way ANOVA, P = 0.09; **Fig. 5A**), indicative of a lack of binding. In contrast, each NTP significantly increased the thermal stability of AbiEii WT (one-way ANOVA, P < 0.0001; **Fig. 5B**), suggesting nucleotide binding is confined to the AbiEii toxin.

**Figure 5.**
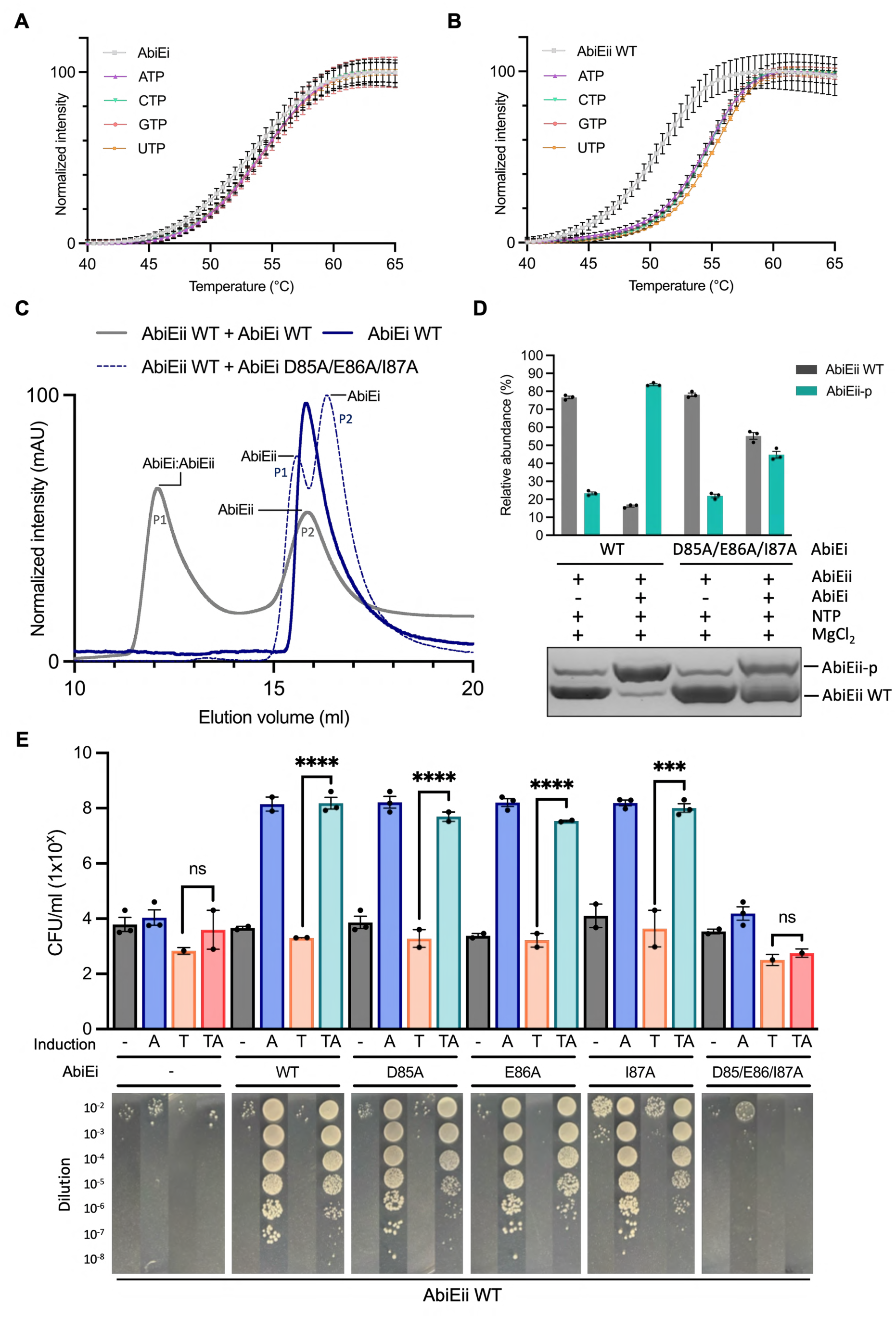
Conserved antitoxin loop residues are essential for antitoxic activity. (**A**, **B**) Thermal shift isotherms of (**A**) AbiEi or (**B**) AbiEii WT incubated in the absence and presence of ATP, CTP, GTP, or UTP. Isotherms are normalized between minima and maxima for presentation and comparison, cropped to the appropriate scale. (**C**) Overlaid SEC traces corresponding to AbiEii WT incubated in the absence and presence of either AbiEi WT or AbiEi D85A/E86A/I87A, and AbiEi WT incubated alone. Samples were analyzed using an analytical Superdex™ 75 increase 10/300 GL SEC column. Chromatograms are normalized between 0-100 for presentation and comparison, cropped to the appropriate scale. (**D**) Phos-Tag SDS-PAGE and densitometric analysis of AbiEii WT incubated with NTP and MgCl_2_, either in the absence or presence of AbiEi WT or AbiEi D85A/E86A/I87A. (**E**) Endpoint viable count antitoxicity assays of *E. coli* DH5α transformed with pTRB482 (AbiEii WT) and either pTA100 empty vector (-), pTRB481 (AbiEi WT), pTRB738 (AbiEi D85A), pTRB739 (AbiEi E86A), pTRB740 (AbiEi I87A), or pTRB741 (AbiEi D85A/E86A/I87A). Overnight cultures were re-seeded into fresh LB supplemented with Ap, Sp and D-glu and grown to mid-log phase. Samples were serially diluted 10^−2^–10^−9^ and spotted onto M9A plates containing Ap and Sp, with or without D-glu, L-ara and IPTG for repression of toxin expression, induction of toxin expression, and induction of antitoxin expression, respectively. Plates were incubated at 37 °C for 48 h, after which they were imaged and colonies counted to determine CFU/ml (one-way ANOVA, P < 0.0001). “A” = antitoxin induced, “T” = toxin induced, “TA” = antitoxin + toxin induced. Assays shown are representative of three independent biological replicates and plotted data represent the mean ± SEM.

As AbiEi did not appear to bind nucleotides, we hypothesized that AbiEi would induce AbiEii auto-phosphorylation. We subsequently re-examined the AbiEi:AbiEii crystal structure to identify antitoxin residues that might play a role in inducing toxin auto-phosphorylation. The overall structure of the AbiEi:AbiEii complex shares good overall similarity to that of the predicted MenA_3_:MenT_3_ AlphaFold model (pTM score = 0.84; RMSD 3.785 Å across 1878 atoms), with respective antitoxins binding across the same face of their cognate toxins (**Supplementary Fig. S5A**). Using molecular dynamics simulations, we previously predicted that the MenA_3_ loop connecting α5-β4 forces the MenT_3_ S78 phosphoacceptor into an inwards-facing conformation that triggers toxin auto-phosphorylation^34^. This flexible loop is also conserved in AbiEi and protrudes into the AbiEii active site in the solved AbiEi:AbiEii structure (**Fig. 3C**). However, in the resolved structure, the AbiEii phosphoacceptor T65 does not reside on a flexible loop, as is the case for MenT_1_ and MenT_3_, but instead comprises the first residue of a central β-strand (**Supplementary Fig. S5B**). Another unique feature of the AbiEi:AbiEii structure is that the flexible AbiEi loop directly interacts with the T65 phosphoacceptor; AbiEi D85 lies a mere 3.1 Å from T65 and forms a hydrogen bond with this residue (**Fig. 3C**), which could suggest a direct role in hydrogen abstraction and activation of T65 for nucleophilic attack of the NTP γ-phosphate. To establish whether D85 is essential for phosphorylation activity, we co-expressed AbiEii WT and AbiEi D85A, then analyzed the resulting sample by Es^+^-ToF MS. To our surprise, AbiEii was still phosphorylated when co-expressed with AbiEi D85A (**Supplementary Fig. S5C**), demonstrating that D85 is not essential for phosphorylation. We therefore reverted to our original hypothesis that antitoxicity is the result of a conformational change to the toxin active site that causes the toxin to self-target, as opposed to direct abstraction of the phosphoacceptor hydroxyl proton by AbiEi.

To investigate whether the unusual positioning of AbiEii T65 in the AbiEi:AbiEii structure was the result of antitoxin-induced conformational changes, we attempted to solve the crystal structure of monomeric AbiEii. However, crystallization efforts were unsuccessful owing to low resolution datasets and poor crystal diffraction. We therefore utilized a predictive AlphaFold model of monomeric AbiEii (AbiEii_apo_; pTM score = 0.92) as a surrogate for alignments onto AbiEii from the AbiEi:AbiEii crystal structure (AbiEii_complex_), allowing us to predict the effects of antitoxin binding on the movement of T65. In the AbiEii_apo_ model, T65 resides on a flexible loop connecting α3-β2 (**Supplementary Fig. S5D**), which matches the position of equivalent phosphoacceptors in MenT_1_ and MenT_3_ (**Supplementary Fig. S5B**). Alignment of AbiEii_apo_ and AbiEii_complex_ structures by Cα atoms returned an RMSD of 0.748 Å (across 201 atoms) and showed a major conformational change in the flexible loop region, causing a 6.8 Å movement of the T65 phosphoacceptor in the AbiEi:AbiEii crystal structure relative to monomeric AbiEii (**Supplementary Fig. S5D**). This suggests that AbiEi antitoxin binding repositions AbiEii T65 closer to the nucleotide-binding pocket. This movement of T65 appears similar to the experimentally observed antitoxin-induced movement of 6.0 Å for MenT_1_ T39 (**Supplementary Fig. S5B**)^34^. The predicted antitoxin-induced movement of T65 supports the hypothesis that a single overarching detoxification mechanism accounts for neutralization of MenT_1_, MenT_3_, and AbiEii.

### Conserved antitoxin loop residues are essential for full antitoxic activity

To verify our hypothesis that AbiEi induces AbiEii auto-phosphorylation we simultaneously mutated the conserved flexible AbiEi looped region to alanine and expressed and purified this triple mutant (AbiEi D85A/E86A/I87A; **Supplementary Fig. S5E**), then assessed its ability to bind AbiEii WT. As we had previously observed (**Fig. 3A**), co-incubation of AbiEii WT and AbiEi WT generated two distinct major peaks during SEC (**Fig. 5C**). Correlation of observed R_st_ values for either peak against calculated R_st_ values for the AbiEi:AbiEii heterodimer and monomeric toxin returned correlation ratios of 1.08 and 0.95, respectively (**Supplementary Fig. S5F**). In contrast, neither of the two peaks captured during SEC following co-incubation of AbiEii WT and AbiEi D85A/E86A/I87A corresponded to the calculated R_st_ of the AbiEi:AbiEii heterodimer (observed/calculated R_st_ ratios of 0.78 and 0.71). However, correlating observed R_st_ values for either peak against calculated values for monomeric AbiEii and AbiEi returned ratios of 0.98 and 0.96 respectively (**Supplementary Fig. S5F**), suggesting that either peak represents monomeric toxin and antitoxin. Together, these data demonstrate that AbiEi residues D85, E86, and I87 are collectively essential for stable complex formation.

Next, we incubated AbiEii WT in the absence and presence of MgCl_2_ and NTP, either with or without AbiEi WT or AbiEi D85A/E86A/I87A mutant, then analyzed the resulting mixtures by Phos-Tag SDS-PAGE. In the absence of antitoxin, weak phosphorylation of AbiEii WT was observed, perhaps the result of abundant substrate under *in vitro* conditions. In contrast, co-incubation with AbiEi WT generated a majority population of AbiEii-p (**Fig. 5D**).

This contrasts the reduced levels of AbiEii-p generated following co-incubation of AbiEii D192A with AbiEi, MgCl_2_, and NTP (**Figs. 1E and F**), supporting the claim that AbiEii D192A exhibits weaker phosphorylation activity than AbiEii WT. In contrast, supplementation with AbiEi D85A/E86A/I87A resulted in a slight increase in the levels of AbiEii-p generated relative to AbiEii WT incubated alone (**Fig. 5D**), but densitometric analysis revealed that AbiEi D85A/E86A/I87A could phosphorylate AbiEii WT with around only half the efficiency of AbiEi WT (**Fig. 5D**). Whilst previous binding assays had indicated that AbiEi D85A/E86A/I87A cannot form a stable heterodimer when mixed with AbiEii WT (**Fig. 5C**), these data suggest that transient complex formation allows some phosphorylation activity, perhaps as has been observed for MenA_3_-MenT_3_^14,34^.

Having now shown that the conserved AbiEi looped region is required for stable TA complex formation and optimal phosphorylation activity *in vitro* (**Figs. 5C and D**), we assessed the global impact of loop mutations on antitoxic activity *in vivo*. We generated a panel of loop residue substitution mutants and tested their ability to neutralize AbiEii toxicity in *E. coli* using endpoint viable count antitoxicity assays. In the absence of antitoxin, AbiEii WT arrested bacterial growth, whereas co-expression of AbiEi WT efficiently blocked toxicity *in vivo* (one-way ANOVA, P < 0.0001; **Fig. 5E**). Similarly, single D85A, E86A, or I87A loop mutations had no effect on the antitoxic activity of AbiEi (one-way ANOVA, P < 0.0001). However, as we had previously observed for MenA_3_^34^, simultaneous mutation of all three loop residues to alanine abolished AbiEi antitoxic activity, resulting in comparable levels of growth inhibition to when AbiEii WT was expressed alone (one-way ANOVA, P = 0.06; **Fig. 5E**). Together, the inability of AbiEi D85A/E86A/I87A to form a stable heterodimeric complex with AbiEii WT (**Fig. 5C**), coupled with the reduced phosphorylation activity of this mutant (**Fig. 5D**), explains the lack of antitoxic activity in *E. coli*. We conclude that, whilst not essential for phosphorylation activity *in vitro*, AbiEi residues D85, E86, and I87 are critical for full antitoxic activity *in vivo*.

### AbiEii exhibits pyrimidine specificity for NTase activity

Having elucidated the mechanism of AbiEii neutralization by AbiEi, we sought to establish the mode of action of the toxin by first re-examining nucleotide specificity. Previous functional characterization demonstrated that AbiEii preferentially binds GTP *in vitro*^20^. However, the observation that AbiEii is phosphorylated when co-expressed with AbiEi indicates that these data are not representative of the nucleotide specificity of the native non-phosphorylated toxin. We subsequently incubated AbiEii WT or AbiEii-p with and without AbiEi, either in the absence or presence of each nucleotide, then analyzed the resulting mixtures by thermal shift. Comparison of mean changes in melting temperature revealed all NTPs provided strong thermal stabilization of AbiEii WT irrespective of the presence of antitoxin (**Fig. 6A**), with UTP providing the greatest stabilization of the four nucleotides in the absence of antitoxin (one-way ANOVA, P < 0.0001). In the presence of AbiEi, both UTP and CTP provided comparable levels of thermal stabilization (**Fig. 6A**), though UTP could still provide stronger thermal stabilization than ATP and GTP (one-way ANOVA, P < 0.05). Conversely, no thermal stabilization of AbiEii-p was observed when incubated with each NTP in the absence of antitoxin, suggesting phosphorylation detrimentally impacts nucleotide binding (**Fig. 6B**). Similarly, only very weak stabilization of AbiEii-p was observed when incubated with each nucleotide in the presence of AbiEi (**Fig. 6B**). Collectively, these data suggest that AbiEii might exhibit preference for UTP as a nucleotide substrate for NTase activity, whilst also providing functional insight into the impact of phosphorylation on nucleotide recognition.

**Figure 6.**
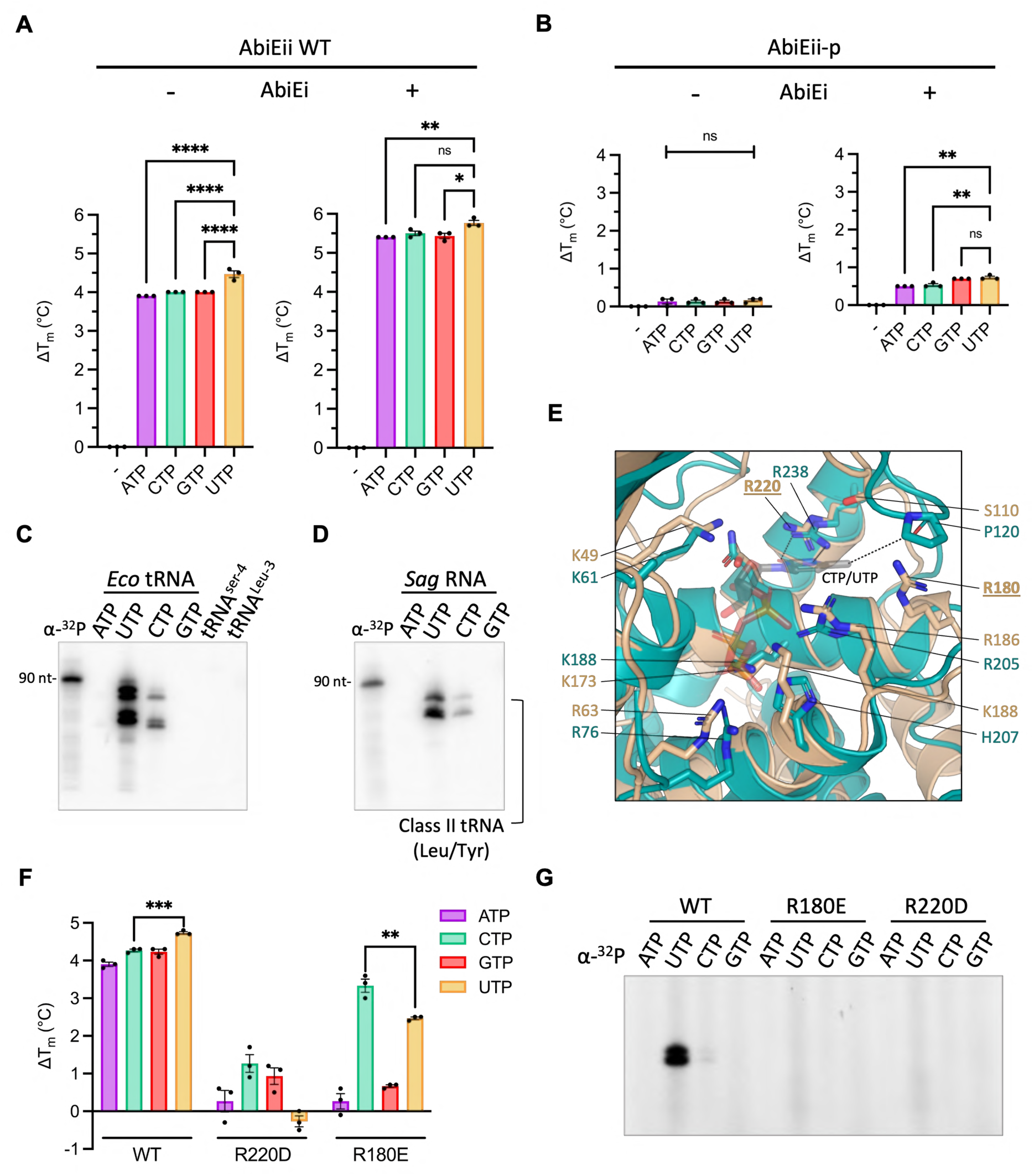
AbiEii exhibits pyrimidine specificity for NTase activity, with a stronger preference for UTP. (**A**, **B**) Mean changes in melting temperature following overnight incubation of either AbiEii WT (**A**) or AbiEii-p (**B**) in the absence and presence of either ATP, CTP, GTP, or UTP, either without (*left*) or with (*right*) AbiEi (one-way ANOVA; **** P < 0.0001, ** P < 0.05). Graphs are representative of three independent biological replicates and plotted data represent the mean ± SEM. (**C**, **D**) Nucleotide transfer assays were performed by incubating 1 µM AbiEii WT in the presence of [α−^32^P] labelled ATP, CTP, GTP, or UTP, and either (**C**) 100 ng purified *E. coli* tRNA mix or 100 ng *in vitro* transcribed tRNA product, or (**D**) 1 µg total RNA from *S. agalactiae*. Reactions were incubated at 37 °C for 10 min, then separated on 6% Urea-PAGE gels and revealed by autoradiography. Gels are representative of two biological replicates. (**E**) Close-up view following superposition of AbiEii (AlphaFold model; pTM score = 0.92) onto MenT_3_ bound to CTP (PDB 8XHR). Structures are shown as cartoon representations colored wheat (AbiEii) and teal (MenT_3_). Residues are shown as sticks with atoms colored red for oxygen, blue for nitrogen, and orange for phosphorous. The pyrimidine backbone of CTP/UTP (based on the MenT_3_:CTP pose) is shaded semi-transparent. (**F**) Mean changes in melting temperature following overnight incubation of either AbiEii WT, AbiEii R220D, or AbiEii R180E in the absence and presence of either ATP, CTP, GTP, or UTP. (**G**) 1 µg total RNA from *S. agalactiae* was incubated with either 1 µM AbiEii WT, AbiEii R220D, or AbiEii R180E in the presence of [α−^32^P] labelled ATP, UTP, GTP, or CTP. Reactions were incubated at 37 °C for 10 min. Assays shown are representative of three independent biological replicates and plotted data represent the mean ± SEM.

Previous characterization of the MenT homologs demonstrated that they modify tRNAs, with only MenT_3_ exhibiting preference for specific isoacceptors^14,23,30^. To examine AbiEii nucleotide specificity, we incubated AbiEii WT in the presence of MgCl_2_, labelled [α−^32^P]-rNTPs, and purified total tRNA from *E. coli*, at 37 °C for 10 min, then visualized tRNAs using autoradiography. As AbiEii shares structural homology with MenT_1_ and MenT_3_ (**Fig. 1B**), we reasoned that it may also target similar tRNAs and so also assessed modification of *in vitro* transcribed *M. tuberculosis* tRNA^Leu-3^ and tRNA^Ser-4^, both of which are targets of MenT_1_ and MenT_3_, respectively^14,23^. In agreement with thermal shift data (**Figs. 6A and B**), tRNA modification assays indicated that AbiEii is pyrimidine-specific for NTase activity, with a preference for UTP over CTP (**Fig. 6C**). Interestingly, AbiEii was unable to modify *M. tuberculosis* tRNA^Leu-3^ or tRNA^Ser-4^, suggesting the toxin exhibits different target specificity than its MenT homologs (**Fig. 6C**). Nucleotide transfer assays were then repeated using total RNA extracted from the native AbiE species *S. agalactiae*, which corroborated preference for UTP and CTP over other NTPs (**Fig. 6D**). Based on the size of modified tRNAs, AbiEii likely exhibits preference for class II tRNAs with long variable arms, reminiscent of MenT_3_^30^.

Like AbiEii, MenT_3_ is also pyrimidine specific for NTase activity and can use both CTP and UTP as substrates, though shows reversed preference of CTP over UTP^14^. As our data indicated that AbiEii exhibits no clear nucleotide preference for *phosphorylation* activity (**Supplementary Fig. S1F**), we speculated that unique toxin-nucleotide interactions are responsible for discriminating between substrates for *NTase* activity. Using the solved crystal structure of MenT_3_ bound to CTP as a guide (PDB 8XHR), we studied the AbiEii active site to identify residues that might govern pyrimidine specificity and preference for UTP over CTP. Superposition of the monomeric AbiEii AlphaFold model onto the MenT_3_:CTP structure returned an RMSD of 3.968 Å (across 1228 atoms). Within the MenT_3_:CTP structure, R238 directly interacts with the O2 and N3 atoms of the cytosine base of CTP^39^; AbiEii R220 is structurally equivalent to MenT_3_ R238 and might also interact with the O2 and N3 atoms of the uracil base of UTP (**Fig. 6E**). We reasoned that mutation of this residue to an aspartate (R220D) might switch the nucleotide preference of the toxin to favor purines over pyrimidines, owing to the presence of N2 and N3 atoms in the adenine/guanine bases of ATP/GTP (**Supplementary Fig. S6A**). Aside from R238, the only other MenT_3_ residue to interact with the cytosine base of CTP is P120, which forms a hydrogen bond from the backbone carbonyl oxygen to the 4-NH_2_ of CTP^39^. This interaction is proposed to favor binding of CTP over UTP, the latter of which lacks the 4-NH_2_ moiety. We therefore hypothesized that the stronger preference of AbiEii for UTP over CTP must be the result of an interaction between the C-4 oxygen of UTP and a different amino acid within the AbiEii structure. Close-up views of the AbiEii active site highlighted the presence of R180 close to where the C-4 carbonyl of UTP is predicted to reside (**Fig. 6E**). We generated a second charge inversion mutant, R180E, which would in theory abolish any interaction between this residue and the C-4 oxygen of UTP, and instead favor interaction with the 4-NH_2_ of CTP (**Supplementary Fig. S6A**).

We subsequently expressed and purified AbiEii R220D and AbiEii R180E in the presence of AbiEi, then analyzed final purified samples by Es^+^-ToF MS. Both mutants retained phosphorylation competency (**Supplementary Fig. S6B**), confirming each could still bind nucleotide(s). However, lower levels of AbiEii-p were generated compared to when AbiEii WT was expressed in the presence of AbiEi (**Supplementary Fig. S1C**), suggesting that R220D and R180E mutations impart a deleterious impact on nucleotide binding, and thus catalytic activity. To test this hypothesis, we first dephosphorylated either mutant using *Sa*STP (**Supplementary Fig. S6C**), then assessed the ability of either mutant to bind nucleotides using thermal shift. ATP, CTP, and GTP provided far weaker thermal stabilization of AbiEii R220D compared to AbiEii WT, whereas UTP slightly reduced thermal stability relative to AbiEii R220D incubated alone (**Fig. 6F**). Surprisingly, purines failed to provide higher levels of thermal stabilization than pyrimidines, suggesting that R220 is not a key determinant of pyrimidine specificity. Thermal stabilization of AbiEii R180E was also far weaker than AbiEii WT when incubated with purines; however, both pyrimidines still provided moderate levels of thermal stabilization relative to AbiEii WT (**Fig. 6F**). Moreover, in line with our hypothesis, CTP now provided significantly higher thermal stabilization than UTP (one-way ANOVA, P = 0.0087), indicating this residue plays a role in determining preference towards UTP or CTP, with AbiEii WT preferring UTP and AbiEii R180E now preferring CTP.

Finally, to establish whether charge inversion mutations had altered nucleotide specificity for NTase activity, we repeated nucleotide transfer assays by co-incubating AbiEii R220D and R180E mutants with total RNA from *S. agalactiae* and each [α−^32^P]-rNTP. However, no modified tRNAs could be detected by autoradiography following incubation with either toxin variant (**Fig. 6G**), suggesting that both are devoid of NTase activity. Thus, whilst thermal shift assays hint at either toxin variant possessing altered catalytic efficiencies and substrate affinities for each nucleotide, the rules for nucleotide usage in NTase catalysis are clearly more nuanced.

### Roles in phage defense and modes of regulation for AbiEi-AbiEii and homologs

AbiEi-AbiEii from *L. lactis* was the first identified two-component AbiE system shown to confer resistance to the 936 family of phages by triggering abortive infection, thereby preventing the release of viable phage progeny from infected cells^7^. To probe whether *S. agalactiae* AbiE functions as an anti-phage system, we performed efficiency of plating (EOP) assays by screening a collection of coliphages against *E. coli* expressing the full AbiEi-AbiEii operon. As a negative control, we also screened against cells expressing an attenuated derivative of the toxin (AbiEii D192A). Of the 32 phages screened, we failed to detect any phenotypical differences in plaque counts or morphology between respective strains (**Supplementary Fig. S7**). Thus, under these conditions, AbiE does not protect *E. coli* from infection.

To identify other AbiEi homologs that might provide phage defense and employ a similar mechanism of antitoxicity, we performed a BLAST search of the entire toxin-antitoxin database^40^ against *S. agalactiae* AbiEi. Of the four experimentally validated antitoxins, only one COG5340 member was annotated as belonging to another family of TA system; RlegA from *Rhizobium leguminosarum*. RlegA shares 22.2% sequence identity with *S. agalactiae* AbiEi and is also encoded downstream from a DUF1814 family member denoted RlegT. Exposure of *E. coli* expressing the RlegTA TA system to phage T7 results in reduced plaque diameter compared to strains expressing RlegA alone^41^, suggesting this system might protect against phage infection. The authors of this study also found that another DUF1814-containing TA system, SanaTA from *Shewanella* sp. ANA-3, could also provide defence against T7 phages lacking gene product (Gp) 4.5 by almost three orders of magnitude^41^. Toxin activation is proposed to result from Lon-mediated degradation of the cognate SanaA antitoxin, with sequestration of Lon by Gp4.5 thought to prevent antitoxin proteolysis in the absence of infection^41^. Whilst the COG5340 domain was not identified in SanaA, InterPro^42^ analysis suggests the antitoxin shares distant homology with members of this superfamily. Despite failing to provide any detectable protection against phage T7^41^, the SdenTA TA system from *Shewanella denitrificans* was also identified as a functional COG2253-COG5340 TA pair. Based on these observations, RlegTA, SanaTA, and SdenTA were scrutinized further to predict whether these systems might also rely on auto-phosphorylation as a mode of toxin regulation.

Antitoxins RlegA and SdenA are both substantially larger than SanaA (205 amino acids and 174 amino acids versus 136 amino acids, respectively), and each possesses additional α-helices and β-sheet bundles at their respective C-termini that are otherwise absent in SanaA (**Fig. 7A**). Superposition of respective antitoxins onto *S. agalactiae* AbiEi (PDB 6Y8Q) revealed shared structural homology between AbiEi and RlegA (RMSD = 3.988 Å across 1077 atoms), SdenA (RMSD = 5.769 Å across 744 atoms), and to a lesser extent SanaA (RMSD = 3.005 Å across 189 atoms; **Fig. 7B**). Closer inspection of each of the superposed structures also revealed that all three AbiEi homologs possess the highly conserved looped region shown to be essential for *S. agalactiae* AbiEi and *M. tuberculosis* MenA_3_ antitoxic activity (**Fig. 7C**)^34^. Akin to AbiEi, each conserved loop is decorated with a quartet of polar and hydrophobic amino acids that might also share similar roles in toxin binding and phosphorylation. Accordingly, predictive AlphaFold models were generated for RlegA:RlegT (pTM score = 0.71), SdenA:SdenT (pTM score = 0.90), and SanaA:SanaT (pTM score = 0.88), working under the assumption these homologous systems would also form stable heterodimeric complexes. Superposition of each of the resulting models onto the AbiEi:AbiEii crystal structure suggests that RlegA:RlegT and SanaA:SanaT complexes, but not that of SdenA:SdenT, share remarkably high similarity to AbiEi:AbiEii (**Figs. 7D and E**). Close-up views also highlight the proximity of flexible antitoxin loops to RlegT and SanaT active site residues, including RlegT T71 and SanaT S65, both of which correspond to AbiEii T65 (**Fig. 7F**). Based on these structural analyses, we predict that RlegA and SanaA antitoxins might employ a similar mechanism of toxin neutralization as AbiEi from *S. agalactiae*. However, further studies are needed to elucidate whether either of these DUF1814 NTases are indeed phosphorylated when expressed in the presence of their cognate antitoxins, as has been demonstrated herein for *S. agalactiae* AbiEi-AbiEii.

**Figure 7.**
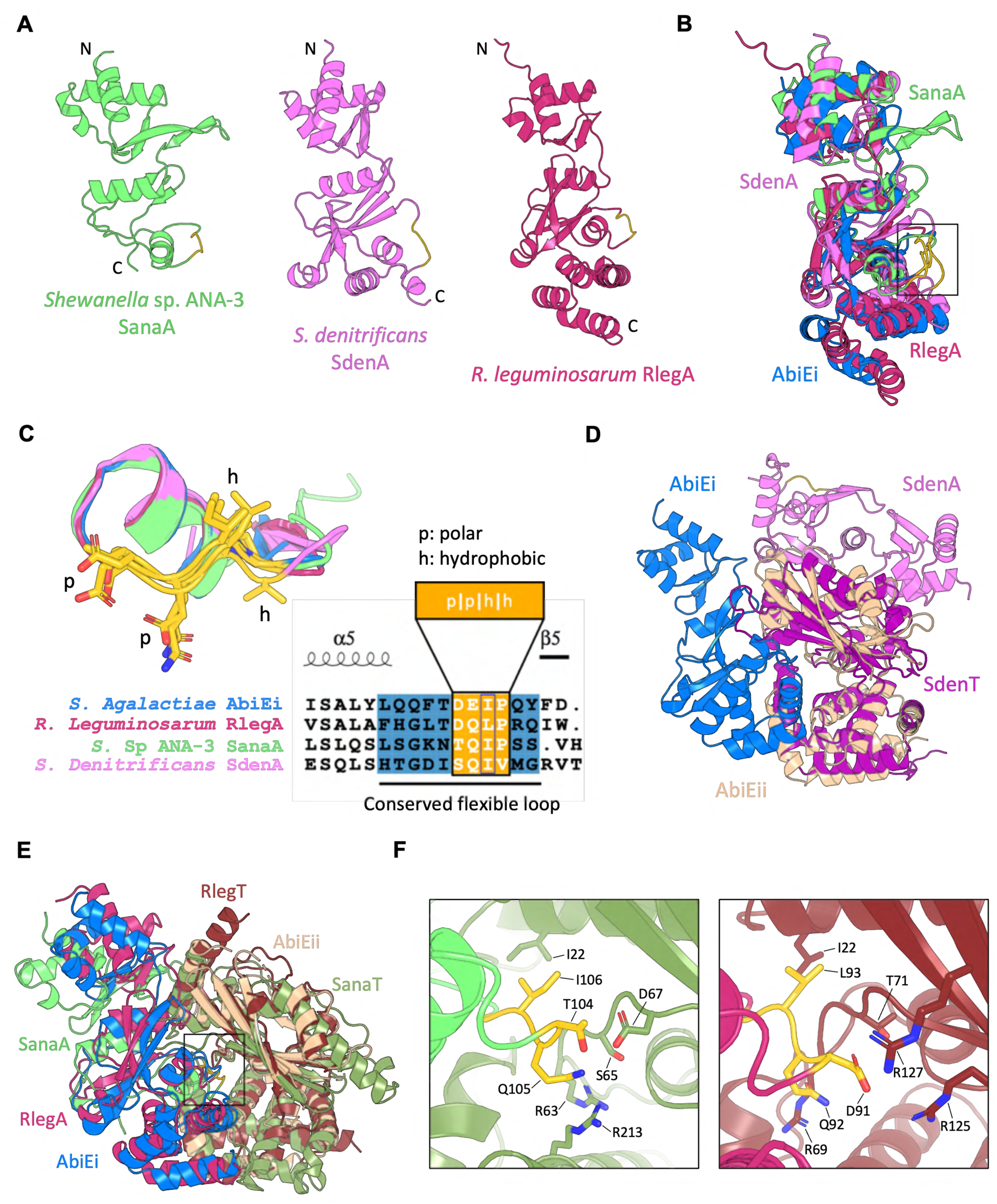
Identification of homologous TA systems that might utilize auto-phosphorylation as a means of toxin neutralization. (**A**) AlphaFold predictive models of *Shewanella* sp. ANA-3 SanaA (pTM score = 0.82), *Shewanella denitrificans* SdenA (pTM score = 0.89), and *Rhizobium leguminosarum* RlegA (pTM score = 0.91). Structures are shown as cartoon representations colored lime green (SanaA), pink (SdenA), and maroon (RlegA). N- and C-termini are indicated. (**B**) Superposition of *S. agalactiae* AbiEi (PDB 6Y8Q) onto SanaA, SdenA, and RlegA AlphaFold models. Structures are shown as cartoon representations colored as in (**A**), with AbiEi colored marine. Conserved antitoxin loop residues are colored gold. (**C**) Close-up view of the boxed region in (**B**) highlighting conservation of essential AbiEi residues in SanaA, SdenA, and RlegA. Residues are shown as sticks with atoms colored red for oxygen and blue for nitrogen. Aligned sequences of respective antitoxin loops (blue background) and amino acids corresponding to AbiEi essential residues (orange background) are also shown. (**D, E**) Superposition of the AbiEi:AbiEii heterodimer onto predictive AlphaFold models of (**D**) SdenA:SdenT (pTM score = 0.90), or (**E**) RlegA:RlegT (pTM score = 0.71) and SanaA:SanaT (pTM score = 0.88). (**F**) Close-up views of the boxed region in (**E**) depicting potential interaction interfaces between SanaA (*left*) or RlegA (*right*) flexible loop residues and respective toxin active site residues, shown as sticks colored red for oxygen and blue for nitrogen.

## Discussion

This study provides biochemical, structural, and biophysical evidence for auto-phosphorylation as a means of toxin neutralization in *S. agalactiae*, a mechanism that had only previously been observed in *M. tuberculosis*. AbiEi binds to AbiEii to form a stable AbiEi:AbiEii heterodimer, leading to subsequent phosphorylation of the toxin in the presence of nucleotide substrates and metal co-factors. Toxin phosphorylation, but not complex formation, efficiently blocks AbiEii toxicity *in vivo*, reclassifying AbiEi-AbiEii as a Type VII TA system. Furthermore, mutagenesis studies highlight the importance of a highly conserved antitoxin loop in modes of binding and phosphorylation, showing that loss of conserved residues translates to an abolition of *in vivo* antitoxic activity. Finally, tRNA modification assays identified UTP as the preferred pyrimidine for AbiEii NTase activity, highlighting additional variability in the substrate specificities of DUF1814 NTases.

We had previously hypothesized that antitoxin-induced phosphorylation is far more widespread than first reported for the *M. tuberculosis* MenAT systems^34^, owing to the ubiquity of DUF1814-COG5340 gene pairs across several major lineages of bacteria and archaea^43^. This study provides an additional layer of evidence in support of this hypothesis, demonstrating that antitoxin-induced phosphorylation is likely conserved amongst different species of pathogenic bacteria. DUF1814 superfamily proteins can be further subclassified into COG2253 and COG4849 NTases, both of which are associated with COG4861 and COG5340 transcriptional regulators respectively (STRING protein-association scores of 95% and 99%). AbiEii, MenT_3_, and MenT_4_ are all COG2253 members encoded downstream of COG5340 antitoxins, but MenT_4_ is not phosphorylated in the presence of MenA_4_^34^. In contrast, MenT_1_ belongs to the COG4849 family and is not associated with a COG4861 family protein, despite undergoing phosphorylation in the presence of the cognate MenA_1_ antitoxin^34^. Analysis of the Conserved Domains Database^44^ reveals MenA_1_ does not belong to any identified superfamily of transcriptional regulators, which matches the absence of the wHTH motif in this antitoxin - a hallmark feature of COG4861 and COG5340 representatives. Therefore, not all COG2253-COG5340 and COG4849-COG4861 pairs belong to the Type VII classification of TA system, and the lack of a COG5340 or COG4861 antitoxin partner does not correlate to an absence of toxin phosphorylation. In the case of AbiEi-AbiEii, the overall mechanism of toxin neutralization is closely resemblant of MenA_1_-MenT_1_ in that toxin phosphorylation is preceded by the formation of a stable TA complex. For MenA_3_-MenT_3_, no such stable complex precedes toxin phosphorylation^34^. Akin to MenT_1_ T39^34^, AbiEii T65 is not essential for NTase activity or toxin neutralization *in vitro* (**Figs. 4B-E and Supplementary Fig. S4**), and higher concentrations of antitoxin are also required to block NTase activity in the absence of phosphorylation (**Fig. 4E and Supplementary Figs. S4C and D**). Whilst structural data had suggested that AbiEi binding leads to steric occlusion of the AbiEii active site (**Supplementary Fig. S3D**), the fact that AbiEii T65A remained toxic in the presence of AbiEi in *E. coli* (**Fig. 4F**) suggests that complex formation is not a viable mode of antitoxicity *in vivo*. This directly contrasts the ability of MenA_1_ to block MenT_1_ T39A toxicity in *M. smegmatis*^23^, and instead matches the inability of MenA_3_ to neutralize MenT_3_ S78A toxicity^36^. It therefore remains unclear as to why certain systems appear to be more reliant on stable complex formation than others, and what the biological relevance of such complexes are in the context of cell growth and metabolism.

Despite the absence of an anti-phage phenotype under the experimental conditions tested in this study (**Supplementary Fig. S7**), we cannot rule out the possibility that AbiE provides phage defence in its native host *S. agalactiae*. Furthermore, the precise mechanisms responsible for toxin activation in response to phage infection remain elusive. Theoretically, toxin activation would be reliant on both the regeneration of functionally active AbiEii by cellular phosphatases, and concomitant disruption of AbiEi activity to prevent negative autoregulation and subsequent phosphorylation of novel toxin transcripts. However, in contrast to the antitoxins of several other TA systems^15,16,45^, AbiEi does not appear to be inherently unstable, as evidenced by remarkably high yields of lone antitoxin following overnight recombinant protein expression in *E. coli* and enhanced thermostability relative to AbiEii *in vitro*. These observations challenge the paradigm that antitoxins are small unstable molecules that are preferentially degraded over stable toxins. Considering the relatively complex nature of AbiEi compared to known protease targets^15,16^, the antitoxin therefore might not represent a suitable target for cellular proteases. Alternatively, given AbiEi exhibits high affinity for its native promoter^22^, viral DNA particles might sequester the antitoxin to disrupt negative autoregulation and prevent AbiEii phosphorylation, potentially leading to activation of the toxin in response to phage infection. A recent study showed that *S. suis* AbiEi could bind to specific DNA sequences in the ICE*Ssu*HN105 plasmid with sub-micromolar potency and, in addition to autoregulation, could also negatively regulate expression of the SezA-SezT TA system^46^, highlighting the ability of AbiEi homologs to recognize DNA sequences besides that of the native promoter. Furthermore, *L. lactis* AbiE could only provide robust phage defence when both AbiEi and AbiEii were expressed^7^, supporting the potential role of the AbiEi antitoxin as a sensory molecule required for activation of AbiEii in response to infection.

Like MenT_1_ and MenT_3_, AbiEii is a pyrimidine-specific NTase capable of blocking translation by perturbing tRNA functionality. However, we have shown that AbiEii is unique in that it is the only known DUF1814 NTase that prefers UTP over CTP as a substrate for NTase activity. Whilst mutagenesis studies hinted at AbiEii playing a key role in governing UTP specificity, other structural features dictating nucleotide usage are yet to be identified. Finally, the role of cellular phosphatases in toxin reactivation, and the underlying mechanisms responsible for blocking AbiEi in response to activating triggers, also require further exploration.

## Methods

### Bacterial strains

*E. coli* strains DH5α (Invitrogen), BL21 (λDE3) (Novagen), and ER2566 (NEB) were routinely grown at 37 °C either in 2x YT or LB media supplemented with, when necessary, 100 μg ml^-1^ ampicillin (Ap), 50 μg ml^-1^ spectinomycin (Sp), 1 mM isopropyl-β-D-thiogalactopyranoside (IPTG), 0.2% w/v L-arabinose (L-ara) or 0.2% w/v D-glucose (D-glu).

### Plasmid constructs

Plasmids are described in **Supplementary Table S1**. All cloning was performed by GenScript Biotech Ltd unless stated otherwise.

### Protein expression and purification

To express AbiEi WT or the AbiEi D85A/E86A/I87A mutant derivative, *E. coli* BL21 (λDE3) cells were transformed with plasmids pTRB525 or pTRB742, respectively. AbiEii mutant derivatives T65A and D192A were expressed by transforming *E. coli* ER2566 with plasmids pTRB713 or pTRB714, respectively. For toxin-antitoxin co-expression, *E. coli* ER2566 cells were transformed with either pPF680 (*abiEi*-*abiEii* WT), pTRB769 (*abiEi*-*abiEii* R180E), pTRB770 (*abiEi*-*abiEii* R220D), or pTRB778 (*abiEi* D85A-*abiEii* WT). To express *Sa*STP, *E. coli* ER2566 cells were transformed with pTRB767.

Single colonies were used to inoculate 130 ml 2x YT for overnight growth at 37 °C with 180 rpm shaking. Starter cultures were re-seeded 1:100 v/v into 2 L baffled flasks containing 1 L 2x YT supplemented with the relevant antibiotic(s). Flasks were subsequently incubated at 37 °C until reaching an OD_600_ of 0.6, at which point the relevant inducing agent(s) was added and cultures incubated overnight at 18 °C. Cells were harvested by centrifugation at 4000 rpm for 20 min at 4 °C and pellets were serially resuspended in ice-cold A500 buffer (20 mM Tris-HCl pH 7.9, 500 mM NaCl, 30 mM imidazole, 10% v/v glycerol), then either snap-frozen in liquid N_2_ and stored at -80 °C or immediately carried forward for purification.

To purify recombinantly expressed proteins of interest, resuspended cells were first disrupted by sonication (45% amplitude, 10 s pulse intervals, 5 min) and clarified by centrifugation at 45000 x g for 40 min at 4 °C. Clarified cell lysate was transferred to a chilled glass beaker on ice and applied to a 5 ml HisTrap™ HP column (Cytiva) pre-equilibrated in A500. The HisTrap™ column was then washed with 50 ml A500, followed by 25 ml A100 buffer (20 mM Tris-HCl pH 7.9, 100 mM NaCl, 10 mM imidazole, 10% v/v glycerol), with bound proteins eluted directly onto a pre-equilibrated 5 ml HiTrap™ Q HP column by washing with 25 ml B100 (20 mM Tris-HCl pH 7.9, 500 mM NaCl, 250 mM imidazole, 10% v/v glycerol). The Q column was re-equilibrated in A100 and transferred to an Åkta™ Pure (Cytiva), with target protein eluted by Anion Exchange Chromatography (AEC) using a salt gradient from 100% A100 to 60% C1000 (20 mM Tris-HCl pH 7.9, 1 M NaCl, 10% v/v glycerol). Chromatographic peak fractions were verified to contain target protein(s) by SDS-PAGE, then pooled. Samples were either incubated overnight with human sentrin/SUMO-specific protease 2 (hSENP2) to facilitate cleavage of the His_6_-SUMO tag, or carried forward directly to the Size Exclusion Chromatography (SEC) purification step (outlined below) in instances where proteins were fused to non-cleavable hexahistidine tags. hSENP2-treated samples were applied to a second HisTrap™ HP column pre-equilibrated in low-imidazole A500 (10 mM imidazole), with flow-through containing untagged target protein collected on ice and subsequently concentrated by centrifugation using the appropriate MWCO spin concentrator (Sartorius). Concentrated protein samples were then applied to a HiPrep^™^ 16/60 Sephacryl^®^ S-200 HR column (S-200; Cytiva) pre-equilibrated in sizing buffer (50 mM Tris-HCl pH 7.9, 500 mM KCl, 10% v/v glycerol) and further purified by SEC. Peak chromatographic fractions were verified to contain target protein(s), then pooled and concentrated as before. Final purified samples were either resuspended in A500 for immediate use, or a 1:2 mixture of storage-to-sizing buffer (storage buffer; 50 mM Tris-HCl pH 7.9, 500 mM KCl, 70% v/v glycerol) for cold storage at −80 °C.

### Mass spectrometry

Purified protein samples were buffer exchanged into 10 mM ammonium bicarbonate using the appropriate MWCO spin concentrator and submitted to the Durham University Department of Chemistry Mass Spectrometry facility at a final concentration of 0.5 mg ml^-1^. Samples were analyzed by positive ion electrospray time-of-flight mass spectrometry (ES^+^-ToF MS), with measurements obtained using a Xevo QToF (Waters, UK) mass spectrometer as previously described^34^. Data were deconvoluted to give neutral masses using MassLynx 4.1 and MaxEnt 1 (Waters, UK).

To conduct LC-MS/MS analysis, samples were prepared as described above and submitted to the Durham University in-house Proteomics facility for ProAlanase digests and subsequent analysis.

### Phos-Tag SDS-PAGE

Incubation mixtures comprising combinations of 10 µM protein, 1 mM GTP, and 10 mM MgCl_2_ were made up to 10 µl with nuclease-free water and incubated at room temperature overnight, then directly mixed with 10 µl 2x Phos-Tag loading dye (125 mM Tris-HCl pH 6.8, 10% v/v BME, 0.002% v/v BPB, 4% w/v SDS, 20% v/v glycerol) and boiled at 95 °C for 5 min. Samples were cooled to room temperature and loaded onto Phos-Tag SuperSep™ 12.5% pre-cast acrylamide gels (Wako pure industries) immersed in 1x Tris Glycine running buffer. Electrophoresis was performed at 180 V constant until the dye front reached the end of the gel, at which point gels were removed and stained as described above. Gels were de-stained in water overnight prior to densitometric analysis of band intensity using ImageJ software, with background normalization and subtraction using the rolling-ball method (radius set to 50 pixels).

### In vitro *phosphorylation assay*

Incubation mixtures were prepared as described for Phos-Tag SDS-PAGE, scaled up to 100 µl final volume. Following overnight incubation, samples were buffer exchanged into 10 mM ammonium bicarbonate using the appropriate MWCO Vivaspin concentrator (Sartorius) and submitted for mass spectrometry analysis as described above.

### In vitro *dephosphorylation assay*

Equal volumes of purified AbiEii-p and *Sa*STP were mixed at a 1:1 mole ratio in the presence of 1x phosphatase buffer (200 mM Tris-HCl pH 7.9, 1 M NaCl, 10 mM DTT, 20 mM MnCl_2_) and incubated at 4 °C overnight. The following morning, this mixture was applied to a 5 ml HisTrap column pre-equilibrated in A500, with flow-through containing untagged *Sa*STP subsequently discarded. The column was then washed with 50 ml A500 to remove residual *Sa*STP, followed by 25 ml B500 to elute AbiEii WT bearing a hexahistidine-tag at its C-terminus. The identity and phosphorylation status of AbiEii WT was finally confirmed by Es^+^-ToF MS.

### Analytical Size Exclusion Chromatography

A Superdex™ 75 increase 10/300 GL SEC column (S-75i; Cytiva) connected to an Åkta™ Pure FPLC system (Cytiva) was pre-equilibrated in 1.2 column volumes of analytical SEC buffer (20 mM Tris-HCl pH 7.9, 150 mM NaCl). A 100 µl capillary loop was first washed with 500 µl nuclease-free water, followed by 500 µl analytical SEC buffer before and between each run, with samples loaded onto the pre-equilibrated loop using a 100 μl Hamilton syringe. To capture the AbiEi:AbiEii complex, 500 µl samples containing 50 nmol of each purified protein were directly mixed and incubated at 4 °C overnight. The following day, samples were concentrated 10-fold and applied to the pre-equilibrated capillary loop. Samples were applied to the column by running 1.2 column volumes of Analytical SEC buffer through the capillary loop at flow rate of 0.5 ml min^-1^. In instances where the content of chromatographic peak fractions required verification by SDS-PAGE, 0.5 ml fractionation was performed, and peak fractions were collected in 96-well deep-plate blocks.

Calibration curves were generated by plotting the elution volumes (V_e_) of controls from LMW/HMW calibration kits (GE healthcare) against their respective known molecular weights. Calibrant mixtures were prepared as 2 separate mixtures, Mix A (3 mg ml^-1^ RNase A, Conalbumin, Carbonic Anhydrase, Aprotinin) and Mix B (3 mg ml^-1^ RNase A, Aprotinin, 4 mg ml^-1^ Ovalbumin), made up to a final volume equal to 0.5% geometric column volume. For determination of column void volume (V_o_), 1 mg ml^-1^ Aldolase was applied to the column as above, with elution volume (V_e_) directly proportional to V_o_. Calibration curve linear regression equations were utilized to convert elution volume to molecular weight. Observed V_e_ values were determined using the Peaks function in Unicorn™ 7 (Cytiva) and converted to partitioning coefficients (K_av_) using the following equation:

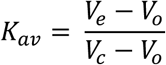

Molecular weight (M_w_) and Stokes radius (R_st_) calibration curves were subsequently plotted using Prism (GraphPad) as K_av_ vs Log_10_(M_w_, kDa) and K_av_ vs Log_10_(R_st_, Å), respectively. Observed R_st_ values were generated by performing linear regression on respective plots using the following equations:

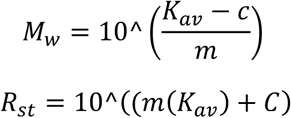

Observed values were then compared against calculated hydrodynamic radii. Radius calculations of inputted crystal structures or AlphaFold predictive models were performed using the HullRad tool (Fluidic Analytics).

### Mass photometry

All mass photometry experiments were performed on the Refeyn TwoMP, using uncoated slides and silicone gaskets. 19µl of analytical SEC buffer was added to the gasket, and the autofocus droplet dilution function was used before loading the protein of interest. Aldolase, Conalbumin, and Ovalbumin were used to generate a standard curve where R^2^ = 1.00 and mass error = 0.6%. Movies were captured for 1 min using the standard image size on Acquire 2024 R1.1. Movies were analyzed using Discover 2024 R2.1. Presented results are the merged analyses of 2 technical replicates for AbiEii D129A, and 4 technical replicates for the AbiEi:AbiEii D192A complex where bin width = 6.

### Protein crystallization and structure determination

The AbiEi:AbiEii complex was concentrated to 15 mg ml^-1^ in Crystal buffer (20 mM Tris-HCl pH 7.9, 150 mM NaCl, 2.5 mM DTT) and crystallization screens were performed using a Mosquito Xtal3 robot (SPT Labtech) using the sitting drop method, with 180:90 nl and 90:90 nl protein-to-condition drops set for each condition screen. AbiEi:AbiEii formed thick cuboidal crystals in condition E6 (0.2 M potassium acetate, 0.1 M MES pH 6.0, 15% v/v pentaerythritol ethoxylate (15/4 EO/OH)) of screen MIDAS*plus*™ (Molecular Dimensions). To harvest crystals for subsequent structural determination, 20 µl screen condition was mixed with 20 µl Cryo buffer (25 mM Tris-HCl pH 7.9, 187.5 mM NaCl, 3.125 mM DTT, 80% glycerol), then added to the protein crystal drop at a 1:1 v/v ratio. Crystals were then immediately extracted from the drop using the appropriately sized nylon loop and transferred to a unipuck immersed in liquid N_2_.

Diffraction data were collected at Diamond Light Source on beamline I04 (**Table 1**). Ten 360° datasets were collected at 0.9795 Å and merged using iSpyB (Diamond Light Source). Data were processed using AIMLESS from CCP4^47,48^ to corroborate spacegroups. The structure was solved by PHASER MR^49^, with the predictive AbiEi:AbiEii AlphaFold structure (pTM score = 0.79) used as a search model. The structure was further built using REFMAC in CCP4^50^, then iteratively refined and built using COOT and Phenix^51,52^ (Ramachandran statistics; 97.12% favored, 2.88% allowed, 0.00% outliers). The quality of the final model was assessed using COOT and the wwPDB validation server^53^. Structural figures, including alignments and superpositions, were generated using PyMol^54^. The APBS electrostatics plugin was used to visualize surface electrostatic charges for models of interest.

### Cell-free protein synthesis

A cell-free transcription/translation coupled assay (PURExpress^®^, Protein synthesis Using Recombinant Elements, NEB) was used to monitor the effects of AbiEii and its derivatives on protein synthesis. 6 µl reactions were assembled on ice and were supplemented with 62.5 ng template DNA (*DHFR*), 0.01-5 µM AbiEii or its variants, 8 U RNase inhibitor, either with or without 0.01-5 µM AbiEi WT. Protein synthesis was performed at 37 °C for 2 h prior to separation of samples on 4-20% SDS-PAGE gradient gels. Gels were subsequently stained overnight with Coomassie Instant Blue (Abcam), then de-stained in water for 2 h and visualized using Image Lab software (Bio-Rad). Densitometric analyses were performed using ImageJ software, with background normalization and subtraction using the rolling-ball method (radius set to 50 pixels).

### Toxicity/antitoxicity assay

To assess the importance of specific amino acids in modes of toxicity and antitoxicity, *E. coli* DH5α cells were co-transformed with either pTRB482 (*abiEii* WT) or pTRB736 (*abiEii* T65A), and either pTA100 empty vector, pTRB481 (*abiEi*), pTRB738 (*abiEi* D85A), pTRB739 (*abiEi* E86A), pTRB740 (*abiEi* I87A), or pTRB741 (*abiEi* D85A/E86A/I87A). Single colonies were used to inoculate 5 ml LB supplemented with D-glu, and starter cultures were grown overnight at 37 °C with 180 rpm shaking. The following day, cultures were re-seeded into fresh LB and grown to mid-log phase. Samples were normalized to an OD_600_ of 1.0 in PBS, then serially diluted and spotted onto M9 agar plates containing either D-glu, L-ara, D-glu and IPTG, or L-ara and IPTG for repression of toxin, induction of toxin, induction of antitoxin, or co-induction of toxin and antitoxin expression, respectively. Plates were incubated at 37 °C for 48 h, then imaged and colonies counted to determine CFU/ml.

### Thermal shift assay

Purified target protein was first labelled with 4 × 10^−3^ µl SYPRO orange dye (Thermo Fisher) per 1 µl protein. Reactions comprising varying concentrations of protein(s) and combinations of 1 mM each NTP were supplemented with 10 µl 2x TSA buffer (20 mM NaH_2_PO_4_, 100 mM NaCl, pH 7.4) and made up to 20 µl with nuclease-free water. Samples were incubated at room temperature overnight in sealed 96-well semi-skirted PCR plates (Starlab), then centrifuged briefly and inserted into a CFX connect real-time qPCR machine (Bio-Rad) for thermal shift analysis. Protein denaturation was performed by incrementally increasing temperature from 25-95 °C. Isotherms were deconvoluted using the NAMI python tool^55^. Melt curves and graphs were generated using Prism (GraphPad).

### In vitro *transcription of tRNAs*

tRNAs were synthesized via *in vitro* transcription using PCR templates that incorporated an integrated T7 RNA polymerase promoter sequence. Primers for *M. tuberculosis* tRNAs are given in **Supplementary Table S1**. T7 TranscriptAid *in vitro* transcription reactions (Thermo Fisher) were carried out in a total volume of 20 µl following the manufacturer’s instructions. Reactions were incubated at 37 °C for 2 h. The resulting tRNA products were extracted using TRIzol™ reagent and stored at a final concentration of 100-200 ng µl^−1^, as determined by NanoDrop analysis.

#### Nucleotide transfer assay

AbiEii WT (or its mutant derivatives) was assayed in 10 µl reaction volumes containing 1x NTase buffer (20 mM Tris-HCl pH 8.0, 10 mM NaCl, 10 mM MgCl_2_) and 1 µCi µl^−1^ of radiolabeled rNTPs [α−^32^P] (Hartmann Analytic) incubated at 37 °C for 10 min. 100 ng *E. coli* tRNA mix, 1 µg *S. agalactiae* total RNA, or 100 ng *in vitro* transcribed tRNA product was used per assay with 1 µM of protein. The 10 µl reactions were purified with Bio-Spin^®^ 6 Columns (Bio-Rad), then mixed with 10 µl of RNA loading dye (95% formamide, 1 mM EDTA, 0.025% SDS, xylene cyanol and bromophenol blue), denatured at 90 °C, and separated on 6% polyacrylamide-urea gels. Gels were vacuum dried at 80 °C and exposed to a phosphorimager screen, then revealed by autoradiography using a Typhoon phosphorimager (GE Healthcare).

#### Phage infection assay

*S. agalactiae* AbiE was screened for phage defence against a panel of coliphages by transforming *E. coli* DH5α with either pTRB773 (*abiEi-abiEii* WT) or pTRB774 (*abiEi-abiEii* D192A). Phage lysate preparation was performed as previously described^57^. For spot test screens, single transformants were used to inoculate 10 ml LB and grown overnight at 37 °C with 180 rpm shaking. 200 µl of each overnight culture were mixed with 3 ml semi-solid ‘top agar’ (0.5% w/v LB agar). The resulting mixtures were poured onto solid LB agar base plates (1.5% w/v LB agar) and dried prior to spotting 3 µl serial dilutions of each coliphage. Plates were incubated overnight at 37 °C prior to imaging.

## Data Availability

The AbiEi:AbiEii crystal structure has been deposited in the Protein Data Bank under accession number 9HLO. All other data needed to evaluate the conclusions in the paper are present in the paper and/or Supplementary Data.

## Acknowledgements

We gratefully acknowledge Diamond Light Source for time on beamline I04 under proposal MX32736. We would like to thank Dr Adrian Brown for performing trypsin and chymotrypsin digests and subsequent LC-MS/MS analyses, and Mr Peter Stokes for performing Es^+^-ToF MS analyses. We also thank Peter Redder and Roland Barriot for useful discussion.

## References

1. Ogura, T. & Hiraga, S. Mini-F plasmid genes that couple host cell division to plasmid proliferation. PNAS 80, 4784–4788 (1983).

2. Kelly, A., Arrowsmith, T. J., Went, S. C. & Blower, T. R. Toxin-antitoxin systems as mediators of phage defence and the implications for abortive infection. Curr Opin Microbiol 73, 102293 (2023).

3. Sonika, S., Singh, S., Mishra, S. & Verma, S. Toxin-antitoxin systems in bacterial pathogenesis. Heliyon 9, e14220 (2023).

4. Hazan, R., Sat, B. & Engelberg-Kulka, H. Escherichia coli mazEF-Mediated Cell Death Is Triggered by Various Stressful Conditions. J Bacteriol 186, 3663–3669 (2004).

5. Fineran, P. C. et al. The phage abortive infection system, ToxIN, functions as a protein-RNA toxin-antitoxin pair. Proc Natl Acad Sci U S A 106, 894–899 (2009).

6. LeRoux, M. et al. The DarTG toxin-antitoxin system provides phage defense by ADP-ribosylating viral DNA. Nat Microbiol 7, 1028 (2022).

7. Garvey, P., Fitzgerald, G. F. & Hill, C. Cloning and DNA sequence analysis of two abortive infection phage resistance determinants from the lactococcal plasmid pNP40. Appl Environ Microbiol 61, 4321–4328 (1995).

8. Tesson, F. et al. Systematic and quantitative view of the antiviral arsenal of prokaryotes. Nat Commun 13, 2561 (2022).

9. Peltier, J. et al. Type I toxin-antitoxin systems contribute to the maintenance of mobile genetic elements in Clostridioides difficile. Commun Biol 3, 1–13 (2020).

10. De Bast, M. S., Mine, N. & Van Melderen, L. Chromosomal toxin-antitoxin systems may act as antiaddiction modules. J Bacteriol 190, 4603–4609 (2008).

11. Bobonis, J. et al. Bacterial retrons encode phage-defending tripartite toxin–antitoxin systems. Nature 609, 144–150 (2022).

12. Bærentsen, R. L. et al. Structural basis for kinase inhibition in the tripartite E. coli HipBST toxin-antitoxin system. Elife 12, 90400 (2023).

13. Yuan, J. et al. Vibrio cholerae ParE2 poisons DNA gyrase via a mechanism distinct from other gyrase inhibitors. Journal of Biological Chemistry 285, 40397–40408 (2010).

14. Cai, Y. et al. A nucleotidyltransferase toxin inhibits growth of Mycobacterium tuberculosis through inactivation of tRNA acceptor stems. Sci. Adv 6, 6651–6680 (2020).

15. Vos, M. R. et al. Degradation of the E. coli antitoxin MqsA by the proteolytic complex ClpXP is regulated by zinc occupancy and oxidation. Journal of Biological Chemistry 298, 101557 (2022).

16. Bordes, P. & Genevaux, P. Control of Toxin-Antitoxin Systems by Proteases in Mycobacterium Tuberculosis. Front Mol Biosci 8, 691399 (2021).

17. Guegler, C. K. & Laub, M. T. Shutoff of host transcription triggers a toxin-antitoxin system to cleave phage RNA and abort infection. Mol Cell 81, 2361–2373 (2021).

18. Beck, I. N. et al. Toxin release by conditional remodelling of ParDE1 from Mycobacterium tuberculosis leads to gyrase inhibition. Nucleic Acids Res 52, 1909–1929 (2024).

19. Qiu, J., Zhai, Y., Wei, M., Zheng, C. & Jiao, X. Toxin–antitoxin systems: Classification, biological roles, and applications. Microbiol Res 264, 127159 (2022).

20. Dy, R. L., Przybilski, R., Semeijn, K., Salmond, G. P. C. & Fineran, P. C. A widespread bacteriophage abortive infection system functions through a Type IV toxin-antitoxin mechanism. Nucleic Acids Res 42, 4590–4605 (2014).

21. Davies, M. R., Shera, J., Van Domselaar, G. H., Sriprakash, K. S. & McMillan, D. J. A Novel Integrative Conjugative Element Mediates Genetic Transfer from Group G Streptococcus to Other ß-Hemolytic Streptococci. J Bacteriol 191, 2257–2265 (2009).

22. Hampton, H. G. et al. AbiEi Binds Cooperatively to the Type IV abiE Toxin-Antitoxin Operator Via a Positively-Charged Surface and Causes DNA Bending and Negative Autoregulation. J Mol Biol 430, 1141–1156 (2018).

23. Xu, X. et al. MenT nucleotidyltransferase toxins extend tRNA acceptor stems and can be inhibited by asymmetrical antitoxin binding. Nat Commun 14, 4644 (2023).

24. Sala, A., Bordes, P. & Genevaux, P. Multiple Toxin-Antitoxin Systems in Mycobacterium tuberculosis. Toxins (Basel) 6, 1002–1020 (2014).

25. Torres, I. et al. Paracoccidioides brasiliensis PbP27 gene: knockdown procedures and functional characterization. FEMS Yeast Res 14, 270–280 (2014).

26. Bassenden, A. V., Park, J., Rodionov, D. & Berghuis, A. M. Revisiting the Catalytic Cycle and Kinetic Mechanism of Aminoglycoside O-Nucleotidyltransferase(2″): A Structural and Kinetic Study. ACS Chem Biol 15, 686–694 (2020).

27. Zhao, Y. et al. Crystal Structure Confirmation of JHP933 as a Nucleotidyltransferase Superfamily Protein from Helicobacter pylori Strain J99. PLoS One 9, e104609 (2014).

28. Tomita, K. & Yamashita, S. Molecular mechanisms of template-independent RNA polymerization by tRNA nucleotidyltransferases. Front Genet 5, 81062 (2014).

29. Aravind, L. & Koonin, E. V. DNA polymerase β-like nucleotidyltransferase superfamily: identification of three new families, classification and evolutionary history. Nucleic Acids Res 27, 1609–1618 (1999).

30. Xu, X. et al. Nucleotidyltransferase toxin MenT extends aminoacyl acceptor ends of serine tRNAs to control Mycobacterium tuberculosis growth. Nat Commun 15, 9596 (2024).

31. Zheng, M., Zheng, M. C., Kim, H. & Lupoli, T. J. Feedback Inhibition of Bacterial Nucleotidyltransferases by Rare Nucleotide l-Sugars Restricts Substrate Promiscuity. J Am Chem Soc 145, 15632–15638 (2023).

32. Chung, C. Z. et al. Gld2 activity is regulated by phosphorylation in the N-terminal domain. RNA Biol 16, 1022 (2019).

33. Wang, X., Yao, J., Sun, Y. C. & Wood, T. K. Type VII Toxin/Antitoxin Classification System for Antitoxins that Enzymatically Neutralize Toxins. Trends Microbiol 29, 388–393 (2021).

34. Arrowsmith, T. J. et al. Inducible auto-phosphorylation regulates a widespread family of nucleotidyltransferase toxins. Nature Communications 2024 15:1 15, 1–20 (2024).

35. Beck, I. N., Usher, B., Hampton, H. G., Fineran, P. C. & Blower, T. R. Antitoxin autoregulation of M. tuberculosis toxin-antitoxin expression through negative cooperativity arising from multiple inverted repeat sequences. Biochem J 477, 2401–2419 (2020).

36. Yu, X. et al. Characterization of a toxin-antitoxin system in Mycobacterium tuberculosis suggests neutralization by phosphorylation as the antitoxicity mechanism. Commun Biol 3, 1–5 (2020).

37. Rantanen, M. K., Lehtiö, L., Rajagopal, L., Rubens, C. E. & Goldman, A. Structure of Streptococcus agalactiae serine/threonine phosphatase: The subdomain conformation is coupled to the binding of a third metal ion. FEBS J 274, 3128 (2007).

38. Krissinel, E. & Henrick, K. Inference of macromolecular assemblies from crystalline state. J Mol Biol 372, 774–797 (2007).

39. Liu, J., Yashiro, Y., Sakaguc hi, Y., omu Suzuki, T. & Tomita, K. Substrate specificity of Mycobacterium tuberculosis tRNA terminal nucleotidyltransferase toxin MenT3. Nucleic Acids Res 2024, 1–15 (2024).

40. Guan, J. et al. TADB 3.0: an updated database of bacterial toxin–antitoxin loci and associated mobile genetic elements. Nucleic Acids Res 52, D784–D790 (2024).

41. Sberro, H. et al. Discovery of Functional Toxin/Antitoxin Systems in Bacteria by Shotgun Cloning. Mol Cell 50, 136–148 (2013).

42. Blum, M. et al. InterPro: the protein sequence classification resource in 2025. Nucleic Acids Res 53, D444–D456 (2025).

43. Makarova, K. S., Wolf, Y. I. & Koonin, E. V. Comparative genomics of defense systems in archaea and bacteria. Nucleic Acids Res 41, 4360–4377 (2013).

44. Wang, J. et al. The conserved domain database in 2023. Nucleic Acids Res 51, D384–D388 (2023).

45. Van Melderen, L., Bernard, P. & Couturier, M. Lon-dependent proteolysis of CcdA is the key control for activation of CcdB in plasmid-free segregant bacteria. Mol Microbiol 11, 1151–1157 (1994).

46. Gu, Q. et al. Type II and IV toxin-antitoxin systems coordinately stabilize the integrative and conjugative element of the ICESa2603 family conferring multiple drug resistance in Streptococcus suis. PLoS Pathog 20, e1012169 (2024).

47. Evans, P. R. & Murshudov, G. N. How good are my data and what is the resolution? Acta Crystallogr. D Biol. Crystallogr. 69, 1204–1214 (2013).

48. Winn, M. D. et al. Overview of the CCP4 suite and current developments. Acta Cryst 67, 235–242 (2011).

49. McCoy, A. J. et al. Phaser crystallographic software. J Appl Crystallogr 40, 658–674 (2007).

50. Murshudov, G. N., Vagin, A. A. & Dodson, E. J. Refinement of macromolecular structures by the maximum-likelihood method. Acta Crystallogr D Biol Crystallogr 53, 240–255 (1997).

51. Emsley, P., Lohkamp, B., Scott, W. G. & Cowtan, K. Features and development of Coot. Acta Crystallogr. D Biol. Crystallogr. 66, 486–501 (2010).

52. Liebschner, D. et al. Macromolecular structure determination using X-rays, neutrons and electrons: recent developments in Phenix. urn:issn:2059-7983 75, 861–877 (2019).

53. Berman, H., Henrick, K., Nakamura, H. & Markley, J. L. The worldwide Protein Data Bank (wwPDB): ensuring a single, uniform archive of PDB data. Nucleic Acids Res 35, D301–D303 (2007).

54. Schrödinger L. The PyMOL Molecular Graphics System, Version 3.0 Schrödinger, LLC.

55. Grøftehauge, M. K., Hajizadeh, N. R., Swann, M. J. & Pohl, E. Protein-ligand interactions investigated by thermal shift assays (TSA) and dual polarization interferometry (DPI). Acta Crystallogr. D Biol. Crystallogr. 71, 36–44 (2015).

56. Fricker, R. et al. A tRNA half modulates translation as stress response in Trypanosoma brucei. Nat Commun 10, 1–12 (2019).

57. Kelly, A. et al. Diverse Durham collection phages demonstrate complex BREX defense responses. Appl Environ Microbiol 89, e0062323 (2023).

