## Supplementary Data for "Antitoxin-induced auto-phosphorylation neutralizes the nucleotidyltransferase toxin AbiEii from *Streptococcus agalactiae* to safeguard global translation"

[d]New England Biolabs, 240 County Road, Ipswich, MA 01938, USA.

†These authors contributed equally.

Keywords: Toxin-antitoxin, phage defence, AbiE, nucleotidyltransferase, phosphorylation

### SUPPLEMENTARY FIGURES

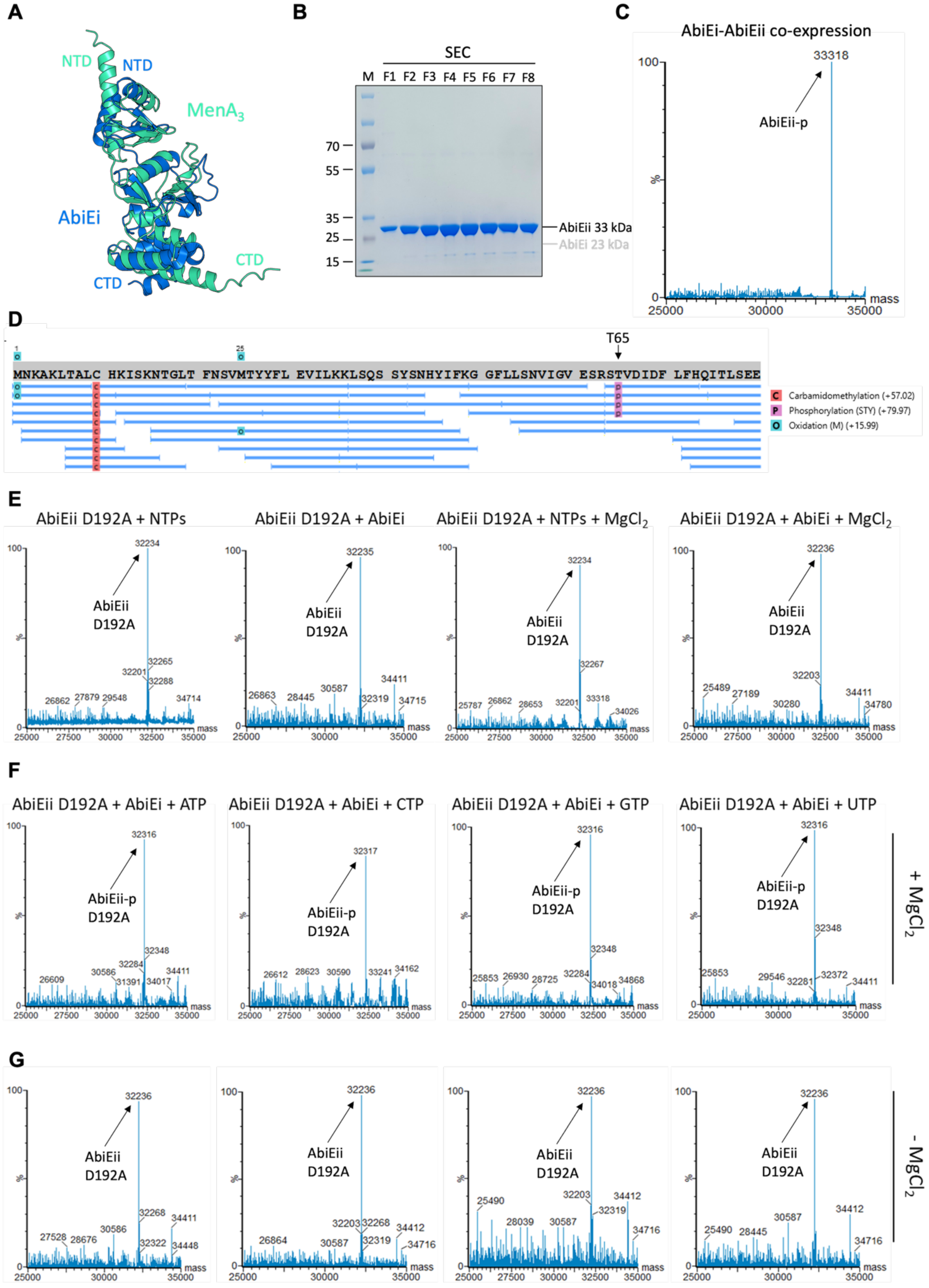


**Supplementary Figure S1.** AbiEii is phosphorylated at T65 in the presence of AbiEi, NTPs, and $MgCl_2$. (**A**) Superposition of AbiEi (PDB 6Y8Q) and $MenA_3$ (AlphaFold model; pTM score = 0.87). Structures are shown as cartoon representations colored marine (AbiEi) and greencyan ($MenA_3$). N- and C-termini are indicated. (**B**) Peak chromatographic fractions following SEC purification of the AbiEi-AbiEii co-expression sample were analyzed by SDS-PAGE. (**C**, **D**) $Es^+$-ToF MS (**C**) and LC-MS/MS (**D**) analysis of the AbiEi-AbiEii co-expression sample confirmed AbiEii phosphorylation in the presence of AbiEi and identified T65 as the phosphoacceptor. (**E**) $Es^+$-ToF MS analysis of AbiEii D192A incubated with combinations of AbiEi, $MgCl_2$, and NTPs. (**F**, **G**) $Es^+$-ToF MS analysis of AbiEii D192A co-incubated with AbiEi and each NTP, either with (**F**) or without (**G**) $MgCl_2$. Mass spectra are representative of three independent biological replicates.

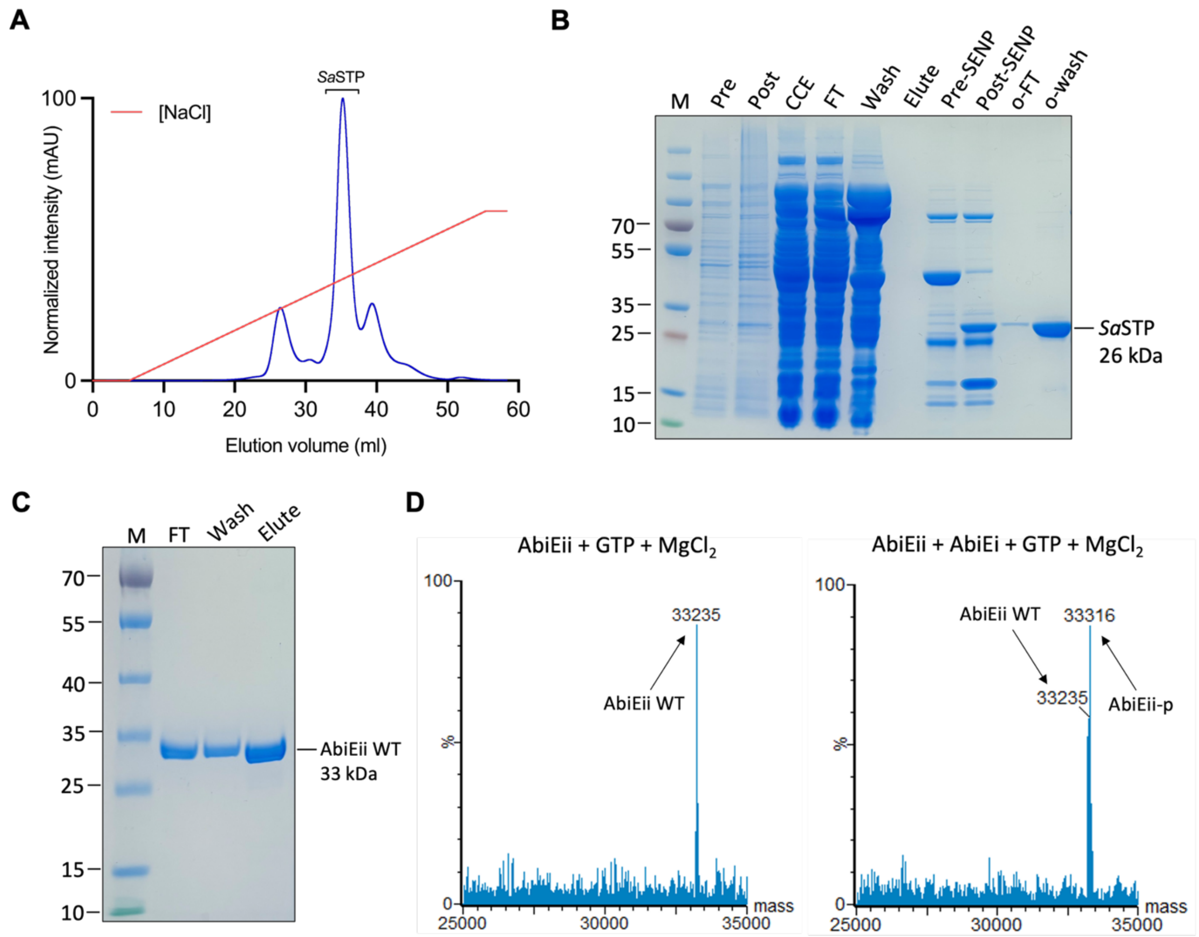


**Supplementary Figure S2.** AbiEii-p dephosphorylation restores phosphorylation competency. (**A**) Chromatographic elution profile of *Sa*STP following Anion Exchange Chromatography. (**B**) *Sa*STP purification fractions were analyzed by SDS-PAGE to verify protein purity. (**C**) Isolation and subsequent purification of non-phosphorylated AbiEii (WT) following dephosphorylation of AbiEii-p. Equal volumes of purified AbiEii-p and *Sa*STP were mixed at a 1:1 mole ratio in the presence of 1x phosphatase buffer (200 mM Tris-HCl pH 7.9, 1 M NaCl, 10 mM DTT, 20 mM $MnCl_2$) and incubated at 4 °C overnight. The following morning, this mixture was applied to a 5 ml HisTrap™ column and washed with 50 ml A500 to elute *Sa*STP, followed by 50 ml B500 to elute $His_6$-tagged AbiEii WT. Purification fractions were analyzed by SDS-PAGE to verify protein purity. (**D**) $Es^+$-ToF MS analysis of purified AbiEii WT incubated with $MgCl_2$ and GTP, either in the absence (*left*) or presence (*right*) of AbiEi. Mass spectra are representative of three independent biological replicates.

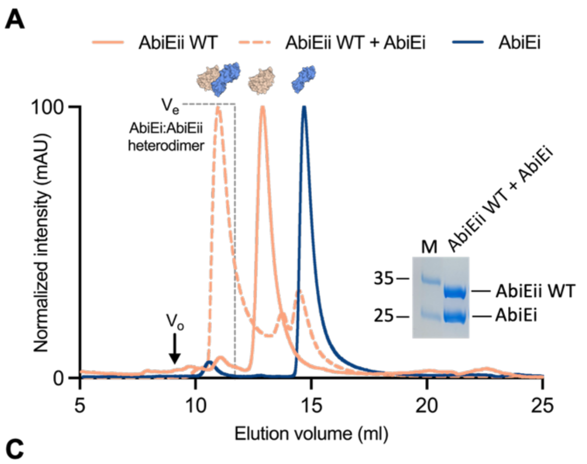


**B**

| Sample | AbiEii WT | AbiEi | AbiEii WT + AbiEi |
|---|---|---|---|
| $V_e$ | 12.90 | 14.69 | 10.99 |
| Observed $M_w$ | 33.7 | 18.4 | 22.9 |
| Calculated $M_w$ | 33.2 | 22.9 | 56.1 |
| Ratio | 1.01 | 0.80 | 1.14 |
| Observed $R_{st}$ | 25.7 | 19.9 | 33.7 |
| Calculated $R_{st}$ | 24.2 | 22.5 | 30.5 |
| Ratio | 1.06 | 0.89 | 1.10 |

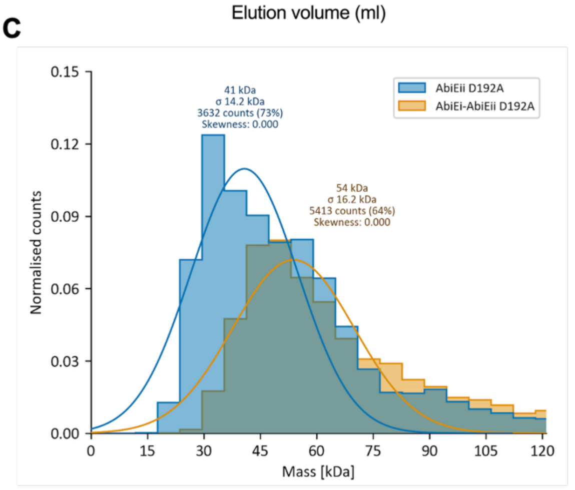


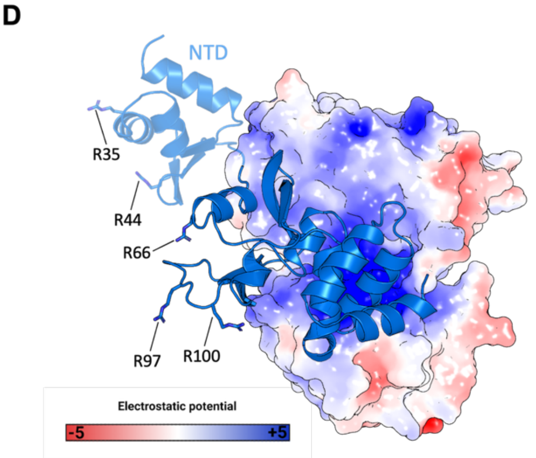


**Supplementary Figure S3.** AbiEi-AbiEii form a stable heterodimer in solution. (**A**) Overlaid SEC traces corresponding to AbiEii WT incubated in the absence and presence of AbiEi, and AbiEi incubated alone. Samples were analyzed using an analytical Superdex™ 75 increase 10/300 GL SEC column. Column void volume ($V_o$) and the calculated elution volume ($V_e$) for the hypothetical AbiEi:AbiEii heterodimer are indicated. Chromatograms are representative of three independent biological replicates and are normalized between 0-100 for presentation and comparison, cropped to the appropriate scale. Peak chromatographic fractions corresponding to the major SEC species formed following AbiEii WT + AbiEi co-incubation were analyzed by SDS-PAGE, revealing the presence of both AbiEi and AbiEii. (**B**) Calculated and observed Molecular Weight ($M_w$) and Stokes Radius ($R_{st}$) values of major peaks corresponding to AbiEii WT incubated in the absence and presence of AbiEi, and AbiEi incubated alone. Observed/calculated ratios are colored green if ≤ 10 %, yellow if > 10 and ≤ 20 %, and red if > 20 % deviation from calculated values. (**C**) Mass photometry analysis of AbiEii D192A incubated in the absence and presence of AbiEi. Graph shows normalized counts from a merged dataset (3 x 1 min movies) for lone toxin (AbiEii D192A) and toxin-antitoxin (AbiEi-AbiEii D192A) mixtures. Lone toxin = 4972 binding events; toxin-antitoxin = 8463 binding events. (**D**) Surface electrostatics of the AbiEii protomer within the AbiEi:AbiEii heterodimer, rotated to show steric occlusion of the NTase active site by the AbiEi CTD. AbiEi DNA-binding residues are shown as sticks. Electrostatics were generated using default settings for the APBS plugin (PyMol), depicting electrostatic potential from −5 $k$B$T$e$^{-1}$ (red) to +5 $k$B$T$e$^{-1}$ (blue), where e is the electron, $T$ is temperature and $k$B is the Boltzmann constant.

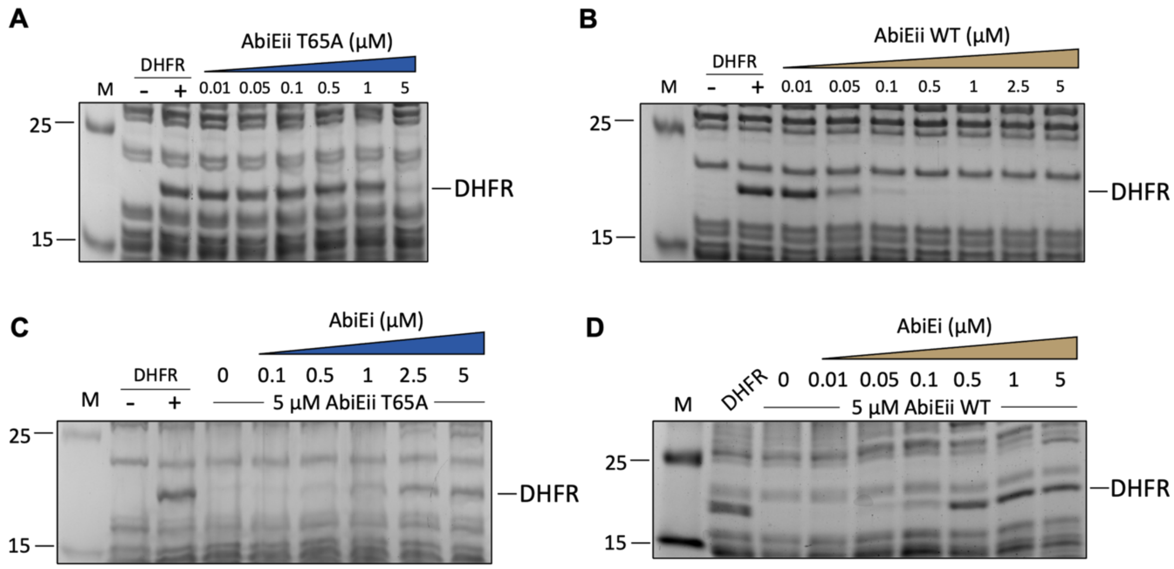


**Supplementary Figure S4.** AbiEi blocks AbiEii T65A NTase activity *in vitro*. (**A**, **B**) *In vitro* transcription/translation reactions assessing levels of DHFR control protein produced in the absence and presence of increasing concentrations of either (**A**) AbiEii WT or (**B**) AbiEii T65A. (**C**, **D**) *In vitro* transcription/translation reactions assessing levels of DHFR control protein produced in the absence and presence of increasing concentrations of AbiEi and either (**C**) AbiEii T65A or (**D**) AbiEii WT (5 µM). Reactions were loaded onto 4-20% gradient gels ran at 180 V for 1 h. Assays shown are representative of three independent biological replicates and plotted data represent the mean ± SEM.

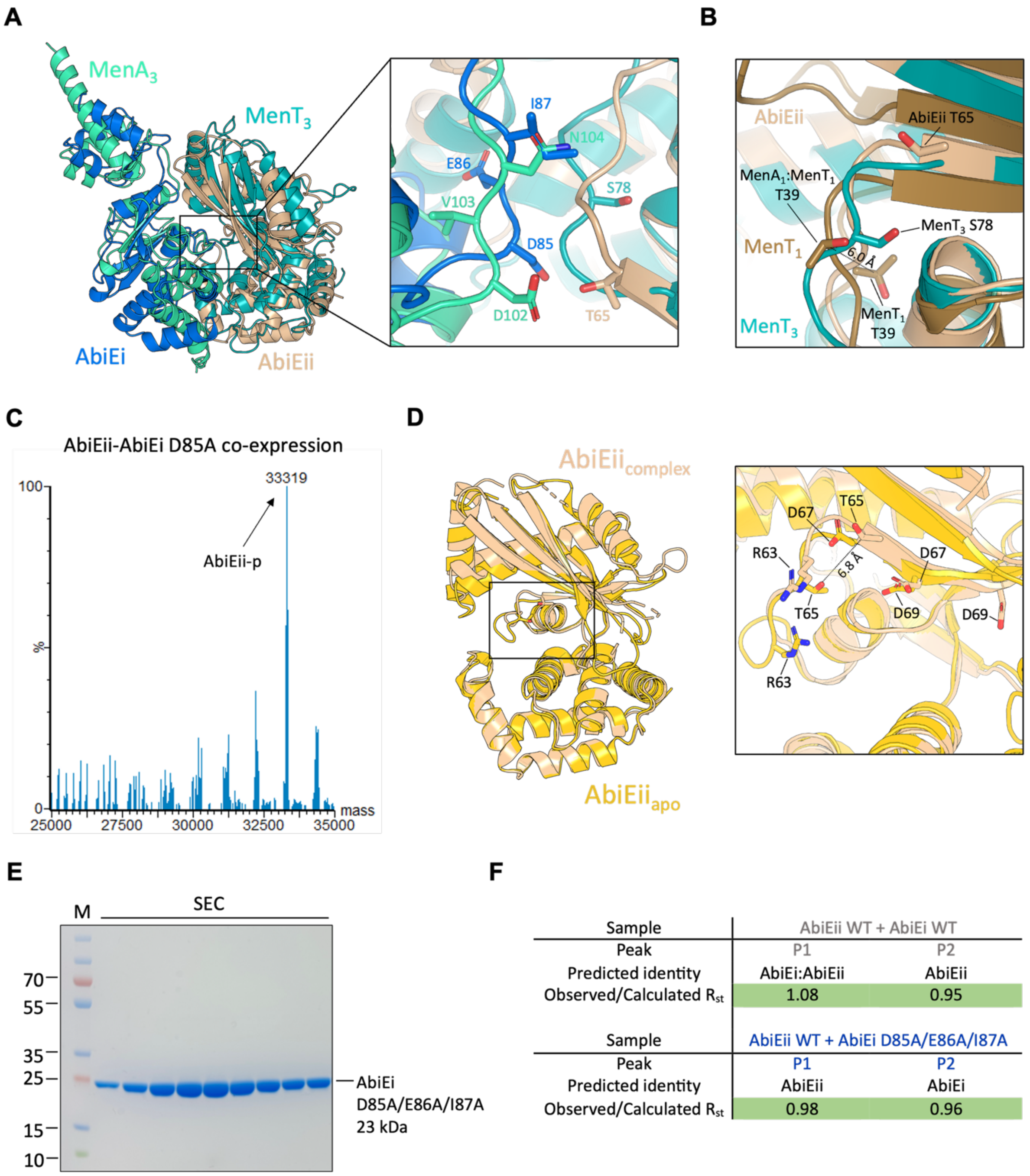


| Sample | AbiEii WT + AbiEi WT | |
|---|---|---|
| Peak | P1 | P2 |
| Predicted identity | AbiEi:AbiEii | AbiEii |
| Observed/Calculated $R_{st}$ | 1.08 | 0.95 |

| Sample | AbiEii WT + AbiEi D85A/E86A/I87A | |
|---|---|---|
| Peak | P1 | P2 |
| Predicted identity | AbiEii | AbiEii |
| Observed/Calculated $R_{st}$ | 0.98 | 0.96 |

**Supplementary Figure S5.** AbiEi does not directly phosphorylate AbiEii. (**A**) Superposition of the AbiEi:AbiEii heterodimer (PDB 9HLO) onto the hypothetical $MenA_3$:$MenT_3$ heterodimer (AlphaFold model; pTM score = 0.84). Structures are shown as cartoon representations colored marine (AbiEi), wheat (AbiEii), greencyan ($MenA_3$), and teal ($MenT_3$). Conserved antitoxin loop residues and cognate toxin phosphoacceptors are shown as sticks with atoms colored red for oxygen and blue for nitrogen. (**B**) Close-up view following Superposition of AbiEii from the AbiEi:AbiEii heterodimer (PDB 9HLO) onto monomeric $MenT_3$ (PDB 8RR6), monomeric $MenT_1$ (PDB 8AN4), and $MenT_1$ from the $MenA_1$:$MenT_1$ heterotrimer (PDB 8AN5). Structures are shown as cartoon representations colored as in (**A**), with $MenT_1$ colored sand. Respective phosphoacceptors are shown as sticks with atoms colored red for oxygen. (**C**) $Es^+$-ToF MS analysis of AbiEii WT expressed in the presence of AbiEi D85A. (**D**) Superposition of monomeric AbiEii ($AbiEii_{apo}$, AlphaFold model; pTM score = 0.92) onto AbiEii from the AbiEi:AbiEii heterodimer ($AbiEii_{complex}$, PDB 9HLO). Structures are shown as cartoon representations colored gold ($AbiEii_{apo}$) and wheat ($AbiEii_{complex}$), with a close-up view highlighting catalytically relevant residues shown as sticks with atoms colored red for oxygen and blue for nitrogen. (**E**) Peak chromatographic fractions following SEC purification of AbiEi D85A/E86A/I87A were analyzed by SDS-PAGE. (**F**) Observed/calculated $R_{st}$ ratios corresponding to each of the major peaks from **Fig. 5C** corresponding to AbiEii WT incubated in the absence and presence of either AbiEi WT or D85A/E86A/I87A mutant. Observed/calculated ratios are colored green if ≤ 10 %, yellow if > 10 and ≤ 20 %, and red if > 20 % deviation from calculated values.

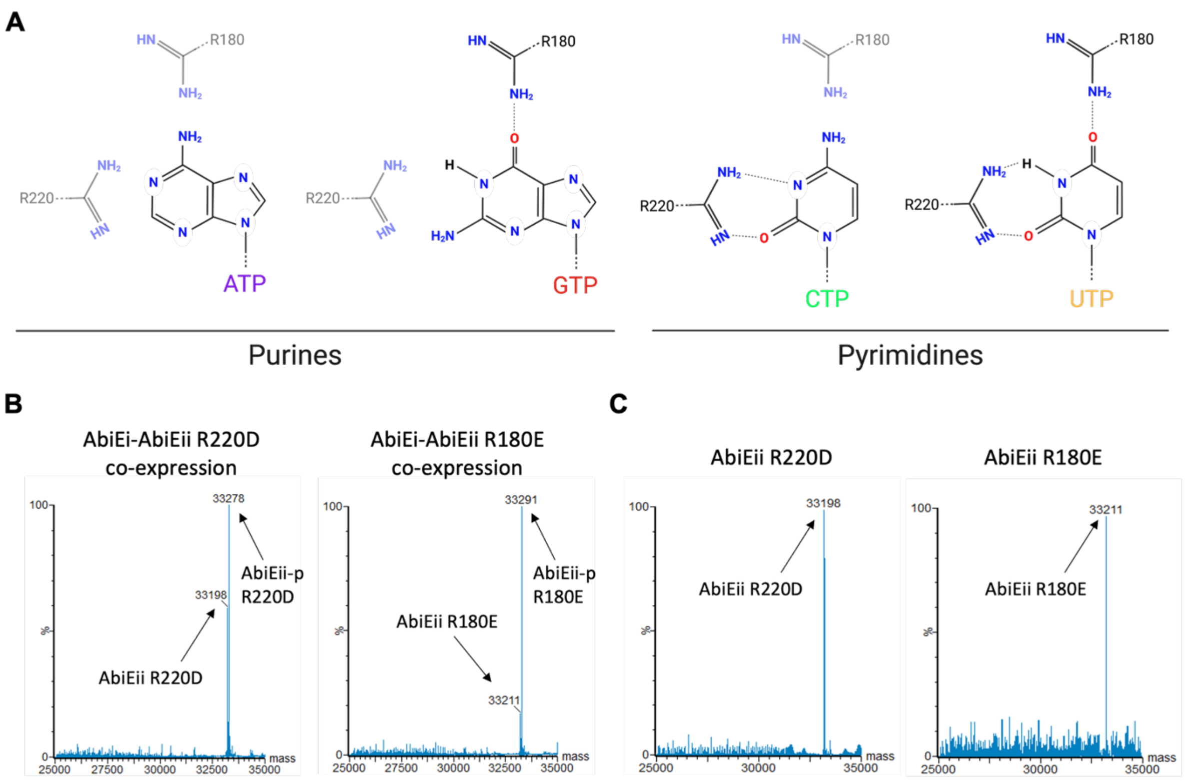


**Supplementary Figure S6.** Design and purification of AbiEii R220D and R180E charge inversion mutants. (**A**) Schematic depicting proposed interactions between AbiEii R180 and R220 and each nucleotide base. (**B**, **C**) Es$^{+}$-ToF MS analysis of AbiEii R220D (*left*) or AbiEii R180E (*right*) expressed in the presence of AbiEi, either before (**B**) or after (**C**) overnight incubation with *Sa*STP.

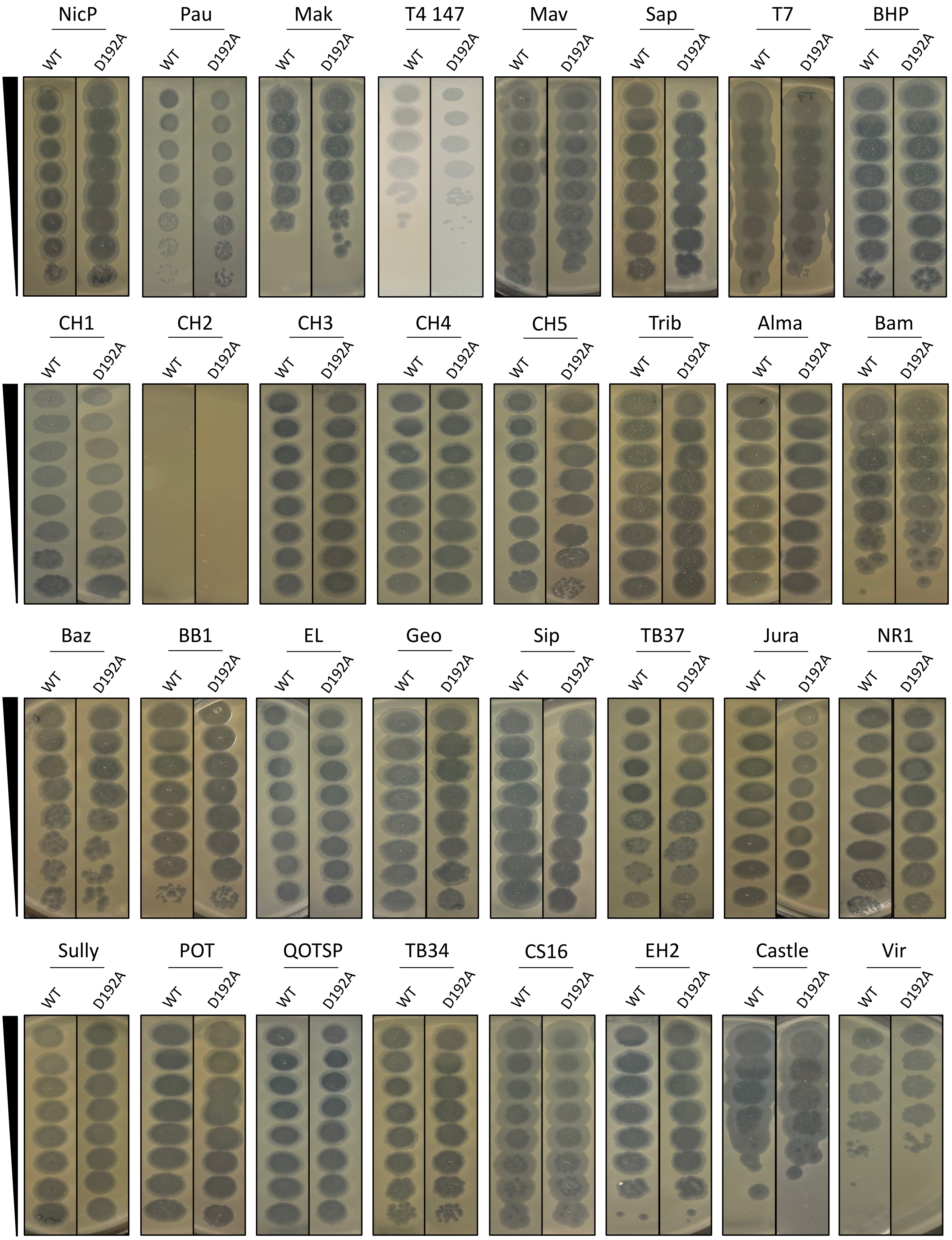


**Supplementary Figure S7.** *S. agalactiae* AbiEi-AbiEii does not provide phage defence against the Durham collection of coliphages. *E. coli* DH5α cells transformed with either pTRB773 (*abiEi-abiEii* WT) or pTRB774 (*abiEi-abiEii* D192A) were screened against the Durham collection of coliphages in phage spot test assays. Single transformants were used to inoculate 10 ml LB and grown overnight at 37 °C with 180 rpm shaking. 200 µl of each overnight culture were mixed with 3 ml semi-solid 'top agar' (0.5% w/v LB agar). The resulting mixtures were poured onto solid LB agar base plates (1.5% w/v LB agar) and dried prior to spotting 3 µl serial dilutions of each coliphage. Plates were imaged after 24 h incubation at 37 °C. Images are representative of three independent biological replicates.

### SUPPLEMENTARY TABLES

**Supplementary Table S1. Plasmids and oligonucleotides used in this study.**

| Vector backbone | Plasmid | Genotype | Source |
|---|---|---|---|
| pBAD30 | pTRB482<br>pTRB736<br>pTRB737<br>pTRB773<br>pTRB774<br>pTRB777 | *abiEii* WT<br>*abiEii* T65A<br>*abiEii* D192A<br>*abiEi-abiEii* WT<br>*abiEi-abiEii* D192A<br>*stp1* | Blower lab<br>GenScript<br>GenScript<br>GenScript<br>GenScript<br>GenScript |
| pQE-80L | pPF680<br>pTRB769<br>pTRB770<br>pTRB778 | $P_{abiE}$; *abiEi*(R35A)-*abiEii*-$His_6$<br>$P_{abiE}$; *abiEi*(R35A)-*abiEii*(R180E)-$His_6$<br>$P_{abiE}$; *abiEi*(R35A)-*abiEii*(R220D)-$His_6$<br>$P_{abiE}$; *abiEi*(R35A/D85A)-*abiEii*-$His_6$ | Fineran lab<br>GenScript<br>GenScript<br>GenScript |
| pSAT1-LIC | pTRB525<br>pTRB742 | $His_6$-SUMO-*abiEi*<br>$His_6$-SUMO-*abiEi* D85A/E86A/I87A | Blower lab<br>GenScript |
| pTA100 | pTRB481<br>pTRB738<br>pTRB739<br>pTRB740<br>pTRB741 | *abiEi*<br>*abiEi* D85A<br>*abiEi* E86A<br>*abiEi* I87A<br>*abiEi* D85A/E86A/I87A | Blower lab<br>GenScript<br>GenScript<br>GenScript<br>GenScript |
| pTRB550 | pTRB713<br>pTRB714<br>pTRB767 | $His_6$-SUMO-*abiEii* T65A<br>$His_6$-SUMO-*abiEii* D192A<br>$His_6$-SUMO-*stp1* | GenScript<br>GenScript<br>GenScript |
| Primers used for *in vitro* tRNA synthesis | | | |
| *Mtb* $tRNA^{Leu-3}$ | For: ATTAATACGACTCACTATAGGTCCGAGTGGCGGAATGGCAGACGCGCTA<br>Rev: TGGTGTCCGAGGGGGGACTT | | |
| *Mtb* $tRNA^{Ser-4}$ | For: ATTAATACGACTCACTATAGGGTGGCGTGGCAGAGCGGCCTAAT<br>Rev: TGGCGGTGGCGGAGGGATTT | | |
